# Divergent *C. elegans* toxin alleles are suppressed by distinct mechanisms

**DOI:** 10.1101/2024.04.26.591160

**Authors:** Stefan Zdraljevic, Laura Walter-McNeill, Giancarlo N. Bruni, Joshua S. Bloom, Daniel H.W. Leighton, Heriberto Marquez, Noah Alexander, Leonid Kruglyak

**Author notes:** Corresponding author: Stefan Zdraljevic, Leonid Kruglyak. Co-first authors (Stefan Zdraljevic, Laura Walter-McNeill).

## Abstract

Toxin-antidote elements (TAs) are selfish DNA sequences that bias their transmission to the next generation. TAs typically consist of two linked genes: a toxin and an antidote. The toxin kills progeny that do not inherit the TA, while the antidote counteracts the toxin in progeny that inherit the TA. We previously discovered two TAs in *C. elegans* that follow the canonical TA model of two linked genes: *peel-1/zeel-1* and *sup-35/pha-1*. Here, we report a new TA that exists in three distinct states across the *C. elegans* population. The canonical TA, which is found in isolates from the Hawaiian islands, consists of two genes that encode a maternally deposited toxin (TMRL-1) and a zygotically expressed antidote (AMRL-1). The toxin induces larval lethality in embryos that do not inherit the antidote gene. A second version of the TA has lost the toxin gene but retains a partially functional antidote. Most *C. elegans* isolates, including the standard laboratory strain N2, carry a highly divergent allele of the toxin that has retained its activity, but have lost the antidote through pseudogenization. Multiple lines of evidence suggest that the N2 *tmrl-1* allele is recognized by piRNAs, leading to MUT-16-dependent 22G siRNA production and post-transcriptional silencing of the transcript. The N2 haplotype represents the first naturally occurring unlinked toxin-antidote system where the toxin is post-transcriptionally suppressed by endogenous small RNA pathways.

## Main Text

Toxin-antitoxin or toxin-antidote (TA) elements are extreme examples of selfish genetic elements that typically consist of two linked genes encoding a toxin and a cognate antidote. The toxin kills individuals that don’t inherit the element and hence lack the antidote to counteract the effects of the toxin (*1–5*). TA elements are ubiquitous in bacteria and have been shown to function as defense mechanisms against bacteriophages, either by directly inhibiting the infection cycle of a phage or by targeting host factors to prevent the spread of mature virions (*6*). Like other immune genes involved in pathogen recognition, TA components are poorly conserved across bacteria because they evolve rapidly to maintain a competitive edge against their target phages (*7, 8*).

While the presence of extremely toxic genes in bacteria can be explained by their role in phage-defense systems, the maintenance of TA elements in metazoans is more mysterious. TA elements are common in hermaphroditic *Caenorhabditis* nematodes (*2–4, 9, 10*), an observation consistent with recent analytical results that selfing can promote the spread of TA elements (*11, 12*). Each of the known *Caenorhabditis* TA elements resides in a hyper-variable genomic region, which suggests that these elements predate the evolution of selfing (*13*) or have contributed to the suppression of gene flow between hyper-variable haplotypes (*11*). TA elements are expected to drive to fixation in outcrossing populations. Once it is fixed or nearly fixed, it loses its selective advantage and there is no selective pressure to maintain it. Therefore, unless a fixed TA provides an additional fitness advantage, the element will likely degrade over time. A recent report suggests that *peel-1*, the toxin component of the first TA element discovered in *C. elegans*, increases host fitness in laboratory conditions, raising the possibility that toxic genes can take on new roles that allow them to be maintained at high frequencies in primarily selfing nematode populations (*14*).

Here, we describe a novel maternally inherited TA element in *C. elegans* with distinctive features. The maternally deposited toxin causes larval arrest rather than embryonic lethality, raising the question of how the toxicity is delayed to this late developmental stage. At the population level, we identified three clades with distinct haplotypes at the new TA locus, the most common of which appears to possess a functional toxin without an antidote. We show that the toxin in this haplotype has been recognized by endogenous piRNA machinery for perpetual silencing by the 22G sRNA pathway. Thus, a vast majority of *C. elegans* strains harbor an unlinked TA system that has no ability to act as a gene drive.

### Identification of a novel *C. elegans* toxin-antidote element

To study the phenotypic effects of genetic variation in *C. elegans*, we generated large cross populations between highly divergent strains—XZ1516 x QX1211 and XZ1516 x DL238. We chose these strains because they are compatible at the two incompatibility loci we previously discovered: *peel-1/zeel-1* and *sup-35/pha-1(2, 3*). We introduced a *fog-2* loss-of-function allele, which feminizes hermaphrodites and prevents them from selfing, into each genetic background to facilitate the construction of large cross populations and intercrossed each population for 10 generations, with minimal selection. Despite minimal selection across each generation, whole-genome sequencing across generations revealed multiple genomic loci with allele frequency distortions, indicating that genetic differences at these loci influenced relative fitness in standard laboratory growth conditions (Fig. S1A). We observed that by generation four of the XZ1516 x QX1211 cross, the XZ1516 allele frequency rose to 75% on the right arm of chromosome V (Fig. 1A). We also observed allele frequency distortion at this region in later generations of the XZ1516 x DL238 cross, which suggested that the same underlying genetic difference was being selected in both crosses. Based on previous studies, we hypothesized that this strong depletion of the QX1211 genotype by generation four is caused by a toxin-antidote (TA) element at this locus (*15*). To test this hypothesis, we performed new crosses between QX1211 and XZ1516 and tracked the phenotypes and genotypes of F2 progeny. We observed that ∼27% of the F2 self-progeny of heterozygous QX1211/XZ1516 F1 hermaphrodites arrested as L1 larvae (Fig. 1B). The observed larval arrest phenotype is reminiscent of the rod-like larval lethal *(rod)* phenotype (Fig. S1B)(*16*). When we crossed QX1211/XZ1516 F1 hermaphrodites to QX1211 males, ∼47% of the progeny exhibited the *rod* phenotype, while we observed no *rod* progeny in the reciprocal cross between QX1211/XZ1516 F1 males and QX1211 hermaphrodites (Fig. 1B). We used PCR genotyping to verify that all *rod* progeny were homozygous for QX1211 alleles at the locus on the right arm of chromosome V that displayed the allele frequency distortion in the mapping populations. This inheritance pattern suggests that the XZ1516 genome encodes a maternally inherited toxin and a linked zygotically expressed antidote that form a novel TA element responsible for the observed allele frequency distortions on the right arm of chromosome V (Fig. 1C). We observed the same F2 phenotypes in crosses between DL238 and XZ1516, indicating that DL238 is a noncarrier of the TA element (Fig. S1C).

**Fig. 1.**
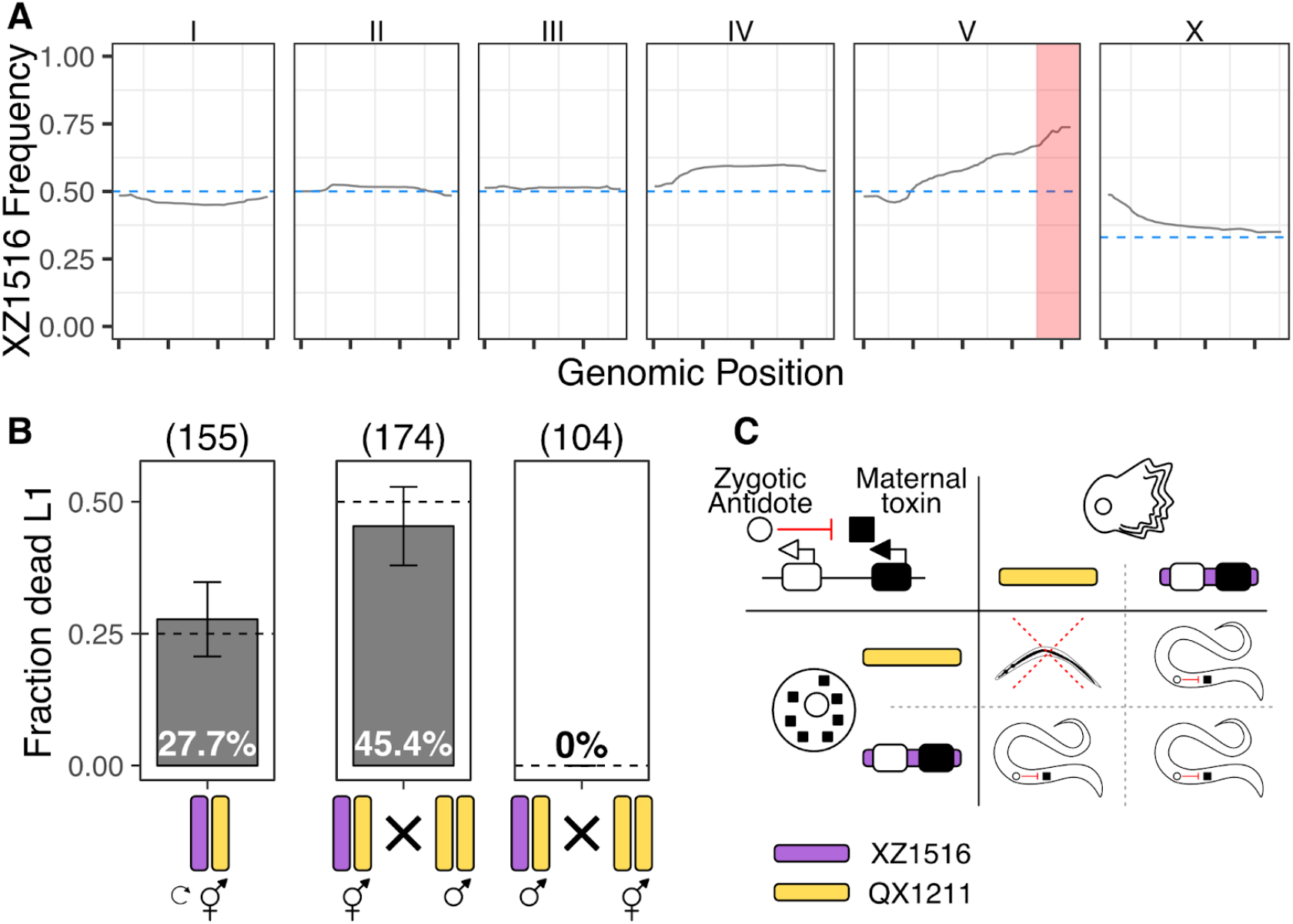
Discovery and characterization of a novel TA. A) The gray line represents the frequency of XZ1516 alleles across the genome after four generations of intercrossing with QX1211. Each panel corresponds to a *C. elegans* chromosome and each x-axis tick indicates 5 Mb. The dotted blue line represents the expected allele frequency for each chromosome with no selection. The region highlighted in red on the right side of chromosome V shows the greatest allele frequency deviation from expectation. B) Crosses between XZ1516 (purple) and QX1211 (yellow) establish the inheritance pattern of the TA element. Bar plots show the fraction of dead L1s observed in each cross. Error bars indicate 95% binomial confidence intervals calculated using the normal approximation method. Crosses from left to right: selfing of XZ1516/QX1211 heterozygous hermaphrodites; XZ1516/QX1211 heterozygous hermaphrodites crossed to QX1211 males; XZ1516/QX1211 heterozygous males crossed to QX1211 hermaphrodites. The observed fraction of dead L1s was not significantly different from the expected fractions for a maternally inherited TA element, exact binomial test. C) Model of the TA inheritance. Punnett square shows the lethality pattern expected in progeny from selfing of XZ1516/QX1211 heterozygous hermaphrodites. A maternally deposited toxin (black square) is present in all progeny and causes L1 lethality unless a zygotically expressed antidote (white circle) is also present.

### Identifying the components of the XZ1516 toxin-antidote element

To isolate the XZ1516 TA element, we introgressed the right arm of chromosome V from XZ1516 into QX1211. We confirmed the identity of the resulting near-isogenic line (NIL) by whole-genome sequencing and verified the presence of the TA element with crosses (Fig. 2A). We were unable to further localize the TA location with standard fine-mapping approaches, likely because of low recombination rates near the ends of *C. elegans* chromosomes (*17*). To overcome the limited natural recombination in this region, we developed a method to induce targeted recombination at double-stranded DNA breaks generated by Cas9 (*18*). This approach enabled us to localize the TA element to a 50 kb region containing 10 candidate genes. We tested these genes for potential toxin or antidote activity by systematically knocking them out in the XZ1516 genetic background (Fig. 2A).

**Fig. 2.**
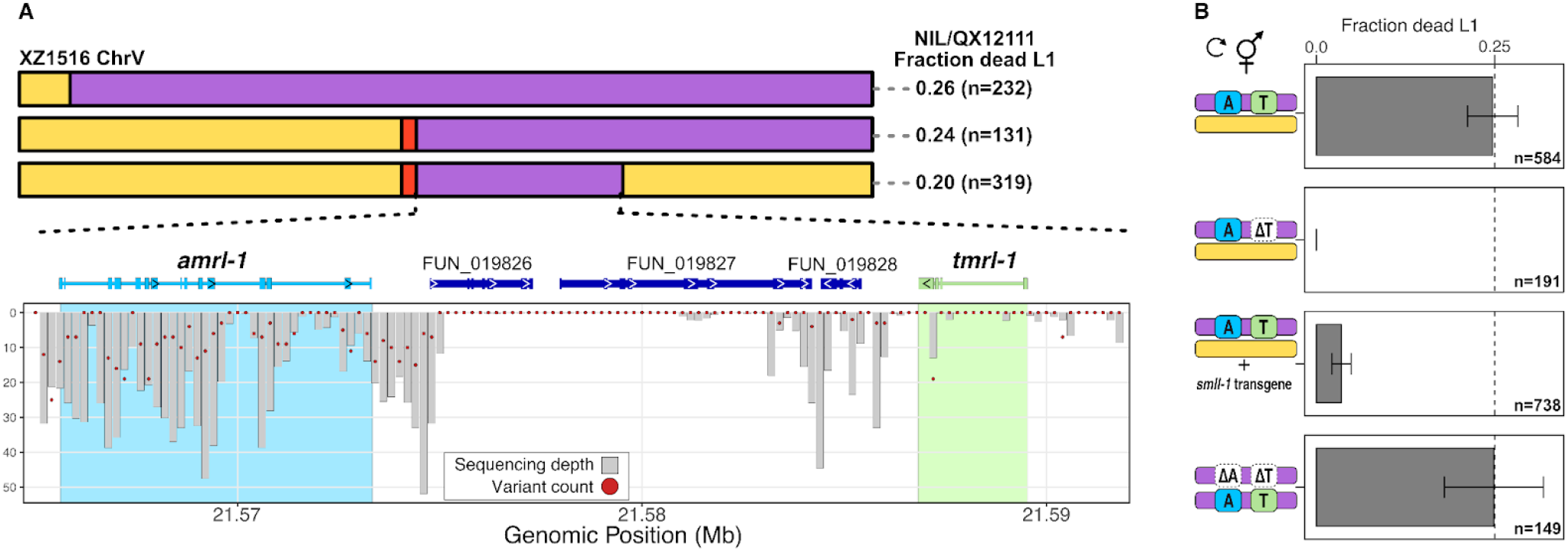
Identification of the TA components. A) Localization of the TA element genes in XZ1516. Top panel: Strain genotypes of near-isogenic lines are displayed as colored rectangles (XZ1516 in purple; QX1211 in yellow; Cas9-induced deletion in red) for chromosome V. The fraction of L1 lethality after selfing of the NIL/QX1211 hermaphrodites is shown to the right of each NIL. The bottom panel depicts a summary of QX1211 sequencing reads aligned to the XZ1516 genome corresponding to the mapped TA element. Gray bars denote short-read sequencing depth in 200 bp windows and red dots denote the number of variants detected between QX1211 and XZ1516 in each window. The XZ1516 and QX1211 genome are so diverged that short reads derived from QX1211 don’t align to the XZ1516 genome in the 200 bp windows with no corresponding read depth, as indicated by a lack of a gray bar. The toxin and antidote genes are highlighted in green and light blue, respectively. B) Knockout and transgenic rescue experiments define the TA components. Bar plots denote the fraction of dead L1s derived from selfing F1 heterozygous individuals. Error bars indicate 95% binomial confidence intervals calculated using the normal approximation method. Blue and green boxes with “A” and “T” indicate intact antidote and toxin genes, respectively; white boxes indicate deletions of these genes. XZ1516 genotypes are depicted in purple and QX1211 genotypes are depicted in yellow. Panels from top to bottom: XZ1516/QX1211 control cross, the observed lethality is not significantly different from the expected 25%; toxin knockout cross to QX1211 the observed lethality is significantly different from the expected 25% p = 1.38e-31; antidote transgenic rescue cross the observed lethality is significantly different from the expected 25% p = 1.22091e-53; toxin and antidote double knockout cross to XZ1516 the observed lethality is not significantly different from the expected 25%. An exact binomial test was used to determine significance.

We isolated three deletion strains that did not induce larval lethality when crossed to QX1211, suggesting that these strains lacked the toxin (Fig. 2B). The computationally predicted gene FUN_019829 is deleted in all three of these strains, and in one of the strains only this gene is deleted, confirming that this gene encodes the toxin. We hereafter refer to FUN_019829 as *tmrl-1* (Toxin-induced Maternal Larval Lethal). We were unable to generate homozygous deletion lines of gene FUN_019825, which suggested that this gene is either essential for survival or encodes the antidote. We successfully isolated homozygous deletion lines of FUN_019825 in a *Δtmrl-1* genetic background, indicating that this gene encodes the antidote. We hereafter refer to FUN_019825 as *amrl-1* (Antidote of Maternal Larval Lethal). We showed that a strain with deletions of both *tmrl-1* and *amrl-1* phenocopies susceptible strains in crosses (Fig. 2B).

To determine whether *amrl-1* is sufficient to suppress *tmrl-1*-induced larval lethality, we constructed a rescue plasmid to drive *amrl-1* expression with a constitutive promoter. We injected the rescue plasmid into XZ1516, crossed individuals harboring the rescue array to a TA-susceptible strain, and selfed the F1 progeny that inherited the array. We observed a dramatic reduction in larval arrest from 25% to 3.5% in F2 progeny, and all F2 progeny that inherited the rescue array survived. These results confirm that *amrl-1* is sufficient to suppress *tmrl-1* toxicity (Fig. 2B).

Long-read RNA sequencing revealed two distinct *tmrl-1* isoforms, a short isoform with three predicted exons and a long isoform with eight predicted exons (Fig. S2A). We constructed plasmids with inducible versions of each *tmrl-1* isoform. When we injected susceptible strains with the short *tmrl-1* isoform array, every F1 individual carrying the array died, with 64% of larvae exhibiting the *rod* phenotype, indicating that uninduced expression levels of the short *tmrl-1* isoform are sufficient to induce lethality. By contrast, we were able to isolate susceptible strains that maintained the long *tmrl-1* isoform array or a short *tmrl-1* isoform array with a premature stop codon in *tmrl-1*. We observed no *rod* progeny upon induction of these arrays, indicating that the short isoform encodes the functional toxin, and that the toxin acts as a protein.

Because lethality only occurs at the L1 stage, we reasoned that *tmrl-1* might be deposited in embryos as a transcript and sequestered from translation. We performed fluorescence *in situ* hybridization (FISH) on developing XZ1516 embryos and larvae with RNA probes that target the *tmrl-1* mRNA. We observed *tmrl-1* puncta as early as the 2-cell embryo stage (Fig. S3A), indicating that *tmrl-1* transcripts are maternally deposited because zygotic transcription does not initiate prior to the 4-cell stage (*19*). At later embryonic and L1 development stages, the *tmrl-1* transcript is localized to two cells that likely correspond to the primordial germ cells (Fig. S3B-C).

### Genomic and population features of the *tmrl-1*/*amrl-1* TA element

The XZ1516 genomic region surrounding the *tmrl-1*/*amrl-1* TA element is hyper-divergent from the reference (N2) genome (*13*). We characterized the genetic variation at this region in the *C. elegans* population by calculating the relatedness of 550 wild isolates (*20*). This analysis separated the population into three distinct clades: an XZ1516-like TA clade which contains 29 strains, a 10-strain clade, and an N2-like susceptible clade composed of 511 strains, including QX1211 and DL238 (Fig. 3A). We verified that the 28 additional isolates with the XZ1516-like haplotype have intact *tmrl-1/amrl-1* genes by aligning sequencing reads from these isolates to the XZ1516 genome assembly. All but four of the isolates in the XZ1516-like clade were collected within three miles of each other on the island of Kauai, two were isolated on Oahu, and one each on Maui and Moloka’i. Four of the isolates from the 10-strain clade were isolated on Maui and the remaining six are globally distributed, while strains with the susceptible haplotype, which represent the majority of the known *C. elegans* isolates, are globally distributed and present on all the Hawaiian islands with the exception of Moloka’i (Fig. 3B).

**Fig. 3.**
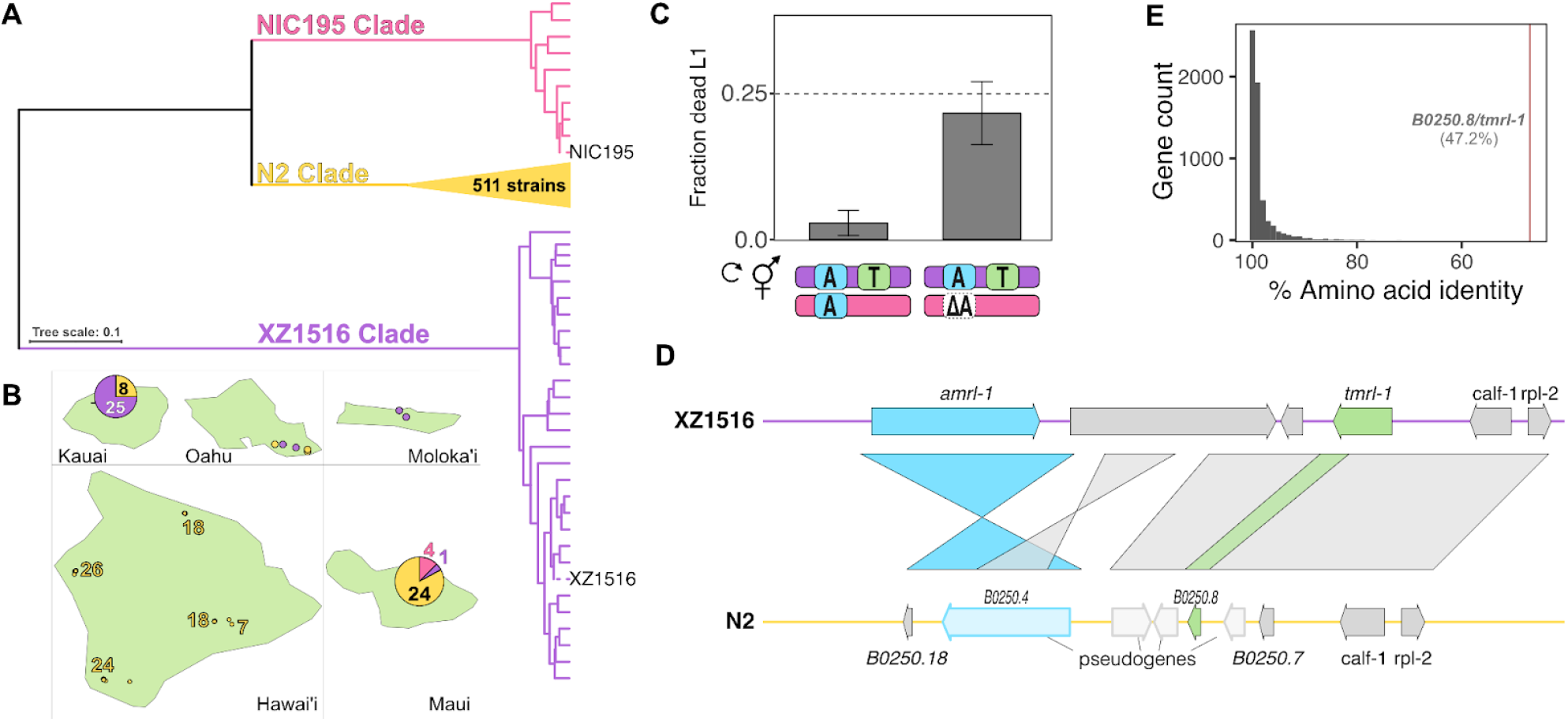
Demographics of the *tmrl-1/amrl-1* TA. A) A dendrogram showing the relatedness of 550 wild *C. elegans* strains at the TA locus. Branches are colored to represent the three distinct clades, where purple denotes the XZ1516-like clade, yellow denotes the N2-like clade, and pink denotes the NIC195-like clade. B) Isolation location of strains collected in Hawaii. Pie charts show the number of isolates from each clade when multiple strains were collected at one location, with colors as in A. C) Bar plots show the fraction of dead L1s in crosses between XZ1516 and NIC195 (left) and between XZ1516 and NIC195 with its antidote allele knocked out (right), indicating that this antidote is active against the XZ1516 toxin. Error bars indicate 95% binomial confidence intervals calculated using the normal approximation method. The observed lethality in the NIC195 x XZ1516 cross is significantly different from the expected 25% *(p* = 6.14e-19, exact binomial test), while the antidote knockout difference is not significantly different. D) Synteny plot of the TA region between the XZ1516 (top) and N2 (bottom) genomes. The TA components *tmrl-1* and *amrl-1* are colored green and blue, respectively. E) Percent amino acid identity of ∼5500 one-to-one orthologs identified between the XZ1516 and N2 genomes. Amino acid identity for *tmrl-1* is indicated with a red line.

The 10-strain clade carries a haplotype that does not contain a gene resembling the toxin. However, this haplotype does carry a divergent *amrl-1* allele that is predicted to contain a full-length coding sequence. We therefore asked whether this *amrl-1* allele is capable of suppressing the toxic effects of *tmrl-1*. We observed the *rod* phenotype in only 3% of F2 progeny derived from crosses between XZ1516 and a representative strain with this haplotype, NIC195, (Fig. 3C), indicating that this antidote is at least partially functional. When we knocked out the *amrl-1* allele in NIC195, 22.5% of F2 progeny were *rod*, confirming that this divergent allele confers reduced susceptibility to the effects of *tmrl-1* (Fig. 3C).

While the previously described *C. elegans* TA elements are characterized by their absence in susceptible strains (*2, 3*), all members of the N2-like susceptible clade harbor a divergent allele of *tmrl-1* with an intact coding sequence, as well as a pseudogenized version of *amrl-1*. The *tmrl-1/amrl-1* genomic region contains several genomic rearrangements between XZ1516 and N2, including likely inversion events that occurred between *amrl-1* and its corresponding divergent N2 allele, *B0250*.*4*; these inversions may have contributed to its pseudogenization (Fig. 3D). While synteny is maintained between *tmrl-1* and the corresponding divergent N2 allele, *B0250*.*8*, many of the surrounding N2 genes are predicted to be pseudogenized. The divergence between *tmrl-1* and *B0250*.*8* is the highest among one-to-one orthologs in the XZ1516 and N2 genomes (nucleotide identity: 63%; protein identity: 47%) (Fig. 3E). We estimated the divergence time for these two alleles under the assumption of neutrality to be between 160 and 325 million generations based on estimates of divergence at synonymous sites (dS)(*21, 22*). This implausibly old estimate suggests that positive selection has been driving the diversification of this gene. The fact that *B0250*.*8* has an intact coding sequence raises the question of whether this gene has maintained its function as a toxin, and if so, how individuals with this haplotype can exist without a functional antidote.

To determine whether *B0250*.*8* acts as a toxin, we used a tetracycline-inducible system to drive the expression of *B0250*.*8* in XZ1516, DL238, and N2. We hatched worms carrying the inducible array on doxycycline plates to induce *B0250*.*8* expression and recorded their phenotypes 48 hours after hatching. All worms expressing *B0250*.*8* displayed a variety of abnormal phenotypes (N2 (n=58); DL238 (n=42); XZ1516(n=61)), which are likely caused by induced expression of *B0250*.*8* in a wide range of tissue types. Notably, we observed the stereotypical *tmrl-1*-dependent *rod* phenotype at low frequencies in all strains (2/58 N2, 5/42 DL238, 4/61 XZ1516). Furthermore, the abnormal phenotypes we observed upon induction of *B0250*.*8* were also seen upon induction of *tmrl-1* (Fig. S4), which suggests that *B0250*.*8* (hereafter N2 *tmrl-1)* has retained its function as a toxin. The presence of a functional toxin and a pseudogenized antidote in N2-like strains suggests that a different mechanism suppresses the toxicity associated with the N2 *tmrl-1* and that this suppression mechanism does not affect the XZ1516 *tmrl-1* toxin (Fig. 1B).

### Small-RNA-mediated suppression of the N2 *tmrl-1* toxin

A potential mechanism that N2-like strains could employ to suppress the activity of *tmrl-1* is RNA interference (RNAi). RNAi pathways are evolutionarily conserved and can act to silence the expression of potentially deleterious genes (*23*). In these pathways, argonaute proteins interact with small RNAs (sRNAs) to transcriptionally and post-transcriptionally regulate gene expression. In *C. elegans*, primary sRNAs initiate the amplification of secondary small interfering RNAs (siRNAs) in perinuclear granules known as *Mutator* foci (*24, 25*). MUT-16 is a glutamine/asparagine (Q/N)-rich protein that is required for the formation of *Mutator* foci at the nuclear periphery of germline nuclei (*25*). Previous work has shown that the N2 *tmrl-1* transcript is heavily targeted by secondary 22G siRNAs, the production of which is dependent on MUT-16 and other *Mutator* foci components (*25*). Animals in which *mut-16* is disrupted with a *mut-16(pk170)* mutation show a 137.7-fold decrease in 22G siRNAs that target *tmrl-1* and a corresponding 23.9-fold increase in the expression level of the gene (*26*) (Fig. S5A-B). Furthermore, high levels of larval arrest occur in mutant strains where *Mutator* foci formation is disrupted, including in *mut-16(pk170)* strains (*27*). Consistent with this report, we observed in a plate-based assay that ∼15% of *Δmut-16* progeny arrested at various larval stages, and 2% of progeny were *rod*, which is suggestive of derepression of *tmrl-1* in N2. We therefore sought to directly test whether *tmrl-1* derepression contributes to larval arrest in the *mut-16(pk170)* strain. To do so, we compared time of flight (TOF) measurements—a proxy for animal length, developmental stage, and growth rate (*28*)—between a strain with a single knockout of *mut-16* and one with a double knockout of *mut-16* and *tmrl-1*(a strain with a single knockout of *tmrl-1* served as a negative control). We observed a reduction in TOF and an increase in the fraction of worms in larval stages in the *mut-16* knockout strain, and these effects were partially rescued in the double knockout strain (Fig. 4). These results indicate that the reduced growth rate observed in the *mut-16* knockout strain is partially mediated by the presence of the N2 *tmrl-1* allele, likely because *tmrl-1* is de-repressed in *mut-16* knockout strains.

**Fig. 4.**
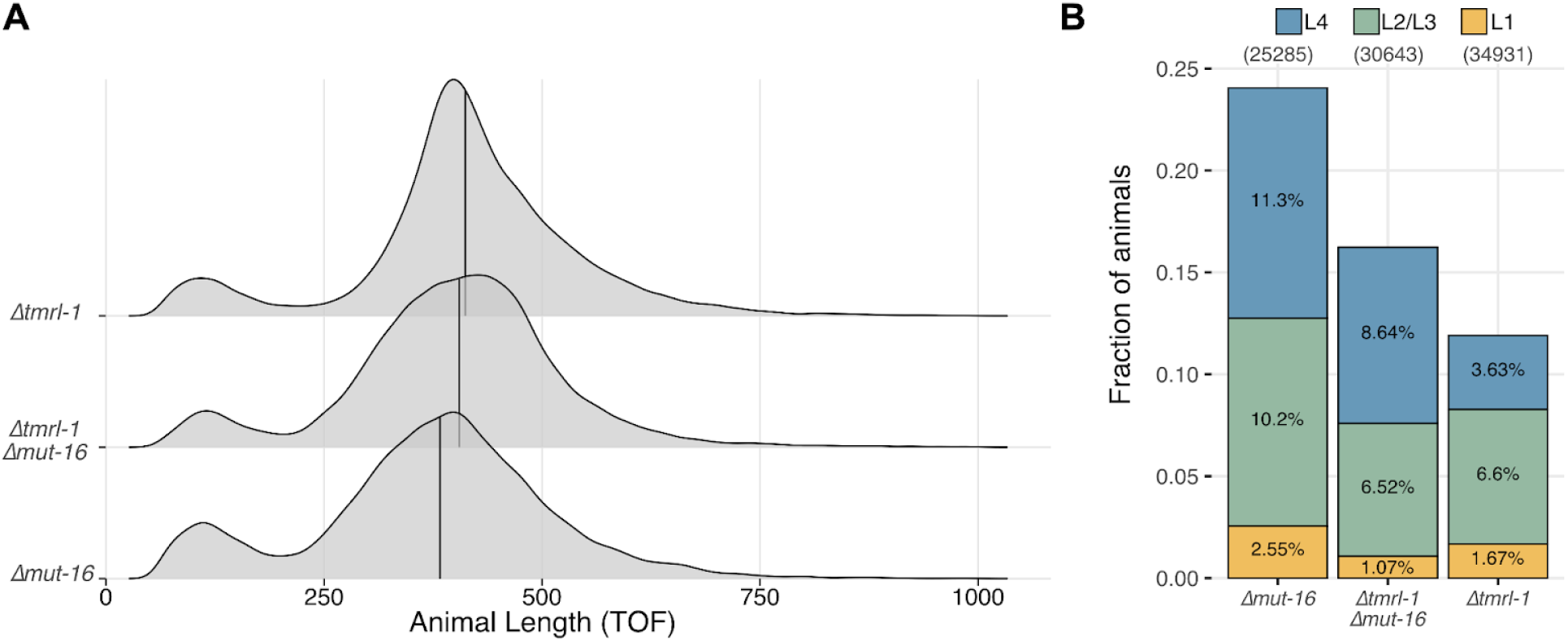
The N2 *tmrl-1* allele contributes to larval arrest in the absence of MUT-16. A) Density plots showing the distribution of animal lengths on the x axis for the *Δtmrl-1, Δmut-16*, and the *Δtmrl-18; Δmut-16* double knockout lines. The distribution of animal lengths are significantly different for all comparisons (Kruskal-Wallis test; *p* = 1.56e-133 for the Δmut-16 to double knockout comparision, *p* = 7.51e-67 for Δ*tmrl-1* to double knockout comparison, and p ≈ 0). B) Animal length data from A) were binned to approximate larval stages as described in the methods. Stacked bar charts of the fraction of animals for each developmental stage for the *Δtmrl-1, Δmut-16, and the Δtmrl-1; Δmut-16* double knockout lines are shown. The fraction of the population is shown on the y axis for each developmental stage – yellow: L1, green: L2/L3, and blue: L4. The fraction of adults is omitted for clarity, but corresponds to the fraction that brings the total to 1 for each genotype.

Amplification of 22G siRNAs can be initiated by different primary sRNAs, including ERGO-1- and ALG-3/4-dependent 26G siRNAs and PRG-1/2-dependent 21U piRNAs. Given that production of MUT-16-dependent 22G siRNAs can be initiated by multiple independent pathways, we queried published sequencing data for sRNAs that are complementary to the N2 *tmrl-1* allele (*29*). This search identified multiple sRNAs that bind throughout the length of the N2 *tmrl-1* transcript. All but one of these sRNAs were not dependent on the argonautes in the queried datasets. We identified one PRG-1-dependent sRNA with a binding site just downstream of two predicted piRNAs, 21ur-8336 and 21ur-14170, which suggests that piRNA recognition of the N2 *tmrl-1* transcript might be involved in its regulation (*30, 31*). In support of this hypothesis, small RNA sequencing of PRG-1-bound piRNAs identified several piRNAs that target the N2 *tmrl-1* transcript (21ur-8336, 21ur-2794, 21ur-2025, 21ur-9583, 21ur-5840, 21ur-4143) (*32, 33*). In line with these observations, 22G siRNAs that target the N2 *tmrl-1* transcript are significantly downregulated in *prg-1(n4357)* gonads as compared to wild type (fold change -17.1; adjusted *p*-value = 2.2e-16) (*26*). The depletion of these PRG-1-dependent siRNAs coincides with a 10.3-fold increase in expression of the N2 *tmrl-1* transcript in *prg-1(n4357)* gonads (*26*). PRG-1-dependent 22G siRNAs produced in the *Mutator* foci interact with the WAGO-1 argonaute in P-granules to silence transcripts (*34*). Recent work has shown that the *N2 tmrl-1* transcript-derived small RNAs co-immunoprecipitated with WAGO-1, providing additional evidence that this transcript is regulated by the endogenous RNAi machinery (*33*) (Fig. S5C). Taken together, these observations suggest that strains with the N2-like haplotype suppress *tmrl-1* toxicity through post-transcriptional silencing mediated by MUT-16-dependent 22G siRNAs that are partially dependent on PRG-1 activity.

## Discussion

We identified a novel toxin-antidote element in *C. elegans* that consists of two genes, *tmrl-1* and *amrl-1*, which encode a maternally deposited toxin and a zygotically expressed antidote, respectively. Unlike the previously characterized *C. elegans* toxins, PEEL-1 and SUP-35, which induce embryonic lethality in susceptible strains, TMRL-1 induces *rod*-like larval lethality. The delayed onset of lethality suggests that the *tmrl-1* transcript is sequestered from translation and degradation throughout embryogenesis and into the early larval stages. This hypothesis is supported by our observations that *tmrl-1* mRNA is distributed across all cells in early embryonic development but is present only in the Z2/Z3 germ cells in older embryos and L1 larvae. While *tmrl-1* has no detectable homology across all sequence databases and only a very low-confidence protein structure prediction, the induction of the *rod* phenotype by TMRL-1 in susceptible strains suggests that it disrupts osmoregulation in the absence of AMRL-1. The *rod* phenotype is caused by fluid filling of the *C. elegans* pseudocoelom and has been observed after laser and genetic ablation of the excretory canal cell, duct cell, pore cell, or CAN neurons (*35–37*), which suggests that these cells are affected by TMRL-1.

A unique feature of the *tmrl-1/amrl-1* element is that three distinct haplotypes of this locus exist across the *C. elegans* population. The XZ1516-like haplotype that we originally identified in two crosses is a canonical toxin-antidote element comprising two linked genes that encode toxin and antidote proteins. The NIC195-like haplotype represents a snapshot of an expected evolutionary trajectory for a toxin-antidote element, in which the toxin is lost through mutation and the antidote is no longer needed to counteract the toxin. This view is supported by the absence of a toxin-like gene in these strains and the accumulation of mutations in the NIC195 version of the antidote that have reduced its ability to counteract the TMRL-1 toxin. These two haplotypes are present in 7% of the known *C. elegans* strains, while the remaining 93% of strains have the N2-like haplotype.

The N2 version of *tmrl-1* is the most divergent one-to-one ortholog between the N2 and XZ1516 genomes. It is important to note that the two orthologs are hyperdivergent at both the nucleotide and the amino acid levels, as indicated by extremely high dN (0.56) and dS (1.77) values and a dN/dS ratio of 0.32. This value of dN/dS is indicative of purifying selection on the protein sequence, in line with our results which show that the N2 version of *tmrl-1* has retained its toxicity. The elevated dN and dS values give implausibly long estimates for the divergence time between these two alleles and suggest that positive selection has been driving the diversification of this gene at the nucleotide level. The absence of an intact version of the antidote gene on this haplotype raised the question of how strains which carry it neutralize the toxin and prompted us to look for an alternative mechanism.

A key difference between the N2 and XZ1516 *tmrl-1* transcripts is the presence of several piRNA binding sites across the N2 transcript. These piRNA binding sites likely enable PRG-1 binding to the N2 *tmrl-1* transcript in P granules before the transcript is shuttled to the *Mutator* foci, where 22G siRNAs are produced by *Mutator* class genes (*38–42*). The 22G siRNAs that target the *N2 tmrl-1* transcript are among the most abundant transcript-specific 22G siRNAs in the N2 genome, and the production of these 22G siRNAs is dependent on both PRG-1 and MUT-16 (*25, 26*). We show that developmental delay phenotypes associated with MUT-16 mutants are partially rescued by removal of the N2 *tmrl-1* gene, suggesting that this gene is likely functional but highly suppressed in wildtype animals by a mechanism that depends on MUT-16. Taken together, our results suggest that most *C. elegans* strains encode a toxic *tmrl-1* gene that is constitutively silenced by unlinked small RNA machinery. While it is impossible to reconstruct the series of events that led to suppression of a toxin by this mechanism, it is likely that small-RNA-mediated suppression of *tmrl-1* arose prior to the loss of the antidote *amrl-1* in the N2 clade. This scenario is reminiscent of the *Stellate* and *Dox* meiotic drive systems in *Drosophila*, in which small-RNA-encoding genes that are unlinked to their target genes are required to downregulate their respective targets to prevent sex distortions in progeny (*43*) and can act as reproductive barriers (*44, 45*). It remains unclear why the N2 *tmrl-1* has not been lost, but we speculate that the divergent *tmrl-1* allele has been maintained in N2-like strains because of a yet-to-be-discovered role it plays in *C. elegans* biology.

## Supporting information

Supplemental Materails

## Author contributions

Conceptualization: SZ

Methodology: SZ, LWM, JSB

Investigation: SZ, LWM, DHWL, HM, NA

Visualization: SZ

Funding acquisition: SZ, GNB, LK

Project administration: SZ, JSB, LK

Supervision: SZ, JSB, LK

Writing – original draft: SZ, LWM, LK

Writing – review & editing: SZ, LWM, LK

## Data and materials availability

All data are available in the manuscript or the supplementary materials.

## References

1. R. W. Beeman, K. S. Friesen, R. E. Denell, Maternal-effect selfish genes in flour beetles. Science 256, 89–92 (1992).

2. E. Ben-David, A. Burga, L. Kruglyak, A maternal-effect selfish genetic element in Caenorhabditis elegans. Science 356, 1051–1055 (2017).

3. H. S. Seidel, M. V. Rockman, L. Kruglyak, Widespread Genetic Incompatibility in \emphC. Elegans Maintained by Balancing Selection. Science 319, 589–594 (2008).

4. E. Ben-David, P. Pliota, S. A. Widen, A. Koreshova, T. Lemus-Vergara, P. Verpukhovskiy, S. Mandali, C. Braendle, A. Burga, L. Kruglyak, Ubiquitous Selfish Toxin-Antidote Elements in Caenorhabditis Species. Curr. Biol. 31, 990–1001.e5 (2021).

5. D. Jurėnas, N. Fraikin, F. Goormaghtigh, L. Van Melderen, Biology and evolution of bacterial toxin–antitoxin systems. Nat. Rev. Microbiol. 20, 335–350 (2022).

6. M. LeRoux, M. T. Laub, Toxin-antitoxin systems as phage defense elements. Annu. Rev. Microbiol. 76, 21–43 (2022).

7. A. J. Shultz, T. B. Sackton, Immune genes are hotspots of shared positive selection across birds and mammals. Elife 8 (2019).

8. M. D. Daugherty, H. S. Malik, Rules of engagement: molecular insights from host-virus arms races. Annu. Rev. Genet. 46, 677–700 (2012).

9. H. S. Seidel, M. Ailion, J. Li, A. van Oudenaarden, M. V. Rockman, L. Kruglyak, A novel sperm-delivered toxin causes late-stage embryo lethality and transmission ratio distortion in C. elegans. PLoS Biol. 9, e1001115 (2011).

10. L. M. Noble, J. Yuen, L. Stevens, N. Moya, R. Persaud, M. Moscatelli, J. L. Jackson, G. Zhang, R. Chitrakar, L. R. Baugh, C. Braendle, E. C. Andersen, H. S. Seidel, M. V. Rockman, Selfing is the safest sex for Caenorhabditis tropicalis. Elife 10 (2021).

11. M. V. Rockman, Parental-effect gene-drive elements under partial selfing, or why do Caenorhabditis genomes have hyperdivergent regions?, bioRxiv (2024) p. 2024.07.23.604817.

12. H. Wang, L. Planche, V. Shchur, R. Nielsen, Selfing Promotes Spread and Introgression of Segregation Distorters in Hermaphroditic Plants. Mol. Biol. Evol. 41 (2024).

13. D. Lee, S. Zdraljevic, L. Stevens, Y. Wang, R. E. Tanny, T. A. Crombie, D. E. Cook, A. K. Webster, R. Chirakar, L. R. Baugh, M. G. Sterken, C. Braendle, M.-A. Félix, M. V. Rockman, E. C. Andersen, Balancing selection maintains hyper-divergent haplotypes in Caenorhabditis elegans. Nat Ecol Evol 5, 794–807 (2021).

14. L. Long, W. Xu, F. Valencia, A. B. Paaby, P. T. McGrath, A toxin-antidote selfish element increases fitness of its host. Elife 12 (2023).

15. A. Burga, E. Ben-David, T. Lemus Vergara, J. Boocock, L. Kruglyak, Fast genetic mapping of complex traits in C. elegans using millions of individuals in bulk. Nat. Commun. 10, 2680 (2019).

16. C. E. Rocheleau, R. M. Howard, A. P. Goldman, M. L. Volk, L. J. Girard, M. V. Sundaram, A lin-45 raf enhancer screen identifies eor-1, eor-2 and unusual alleles of Ras pathway genes in Caenorhabditis elegans. Genetics 161, 121–131 (2002).

17. M. V. Rockman, L. Kruglyak, Recombinational landscape and population genomics of \emphCaenorhabditis elegans. PLoS Genet. 5, e1000419 (2009).

18. S. Zdraljevic, L. Walter-McNeill, H. Marquez, L. Kruglyak, Heritable Cas9-induced nonhomologous recombination in C. elegans. microPublication Biology 2023 (2023).

19. S. Robertson, R. Lin, “Chapter One - The Maternal-to-Zygotic Transition in C. elegans” in Current Topics in Developmental Biology, H. D. Lipshitz, Ed. (Academic Press, 2015; https://www.sciencedirect.com/science/article/pii/S0070215315000290)vol. 113, xpp. 1–42.

20. D. E. Cook, S. Zdraljevic, J. P. Roberts, E. C. Andersen, CeNDR, the Caenorhabditis elegans natural diversity resource. Nucleic Acids Res. 45, D650–D657 (2017).

21. J. H. Gillespie, C. H. Langley, Are evolutionary rates really variable? J. Mol. Evol. 13, 27–34 (1979).

22. C. G. Thomas, W. Wang, R. Jovelin, R. Ghosh, T. Lomasko, Q. Trinh, L. Kruglyak, L. D. Stein, A. D. Cutter, Full-genome evolutionary histories of selfing, splitting, and selection in Caenorhabditis. Genome Res. 25, 667–678 (2015).

23. A. K. Rogers, C. M. Phillips, A Small-RNA-Mediated Feedback Loop Maintains Proper Levels of 22G-RNAs in C. elegans. Cell Rep. 33, 108279 (2020).

24. C. J. Uebel, D. C. Anderson, L. M. Mandarino, K. I. Manage, S. Aynaszyan, C. M. Phillips, Distinct regions of the intrinsically disordered protein MUT-16 mediate assembly of a small RNA amplification complex and promote phase separation of Mutator foci. PLoS Genet. 14, e1007542 (2018).

25. C. M. Phillips, T. A. Montgomery, P. C. Breen, G. Ruvkun, MUT-16 promotes formation of perinuclear mutator foci required for RNA silencing in the C. elegans germline. Genes Dev. 26, 1433–1444 (2012).

26. K. J. Reed, J. M. Svendsen, K. C. Brown, B. E. Montgomery, T. N. Marks, T. Vijayasarathy, D. M. Parker, E. O. Nishimura, D. L. Updike, T. A. Montgomery, Widespread roles for piRNAs and WAGO-class siRNAs in shaping the germline transcriptome of Caenorhabditis elegans. Nucleic Acids Res. 48, 1811–1827 (2020).

27. A. K. Rogers, C. M. Phillips, Disruption of the mutator complex triggers a low penetrance larval arrest phenotype. microPublication Biology, doi: 10.17912/micropub.biology.000252 (2020).

28. E. C. Andersen, T. C. Shimko, J. R. Crissman, R. Ghosh, J. S. Bloom, H. S. Seidel, J. P. Gerke, L. Kruglyak, A Powerful New Quantitative Genetics Platform, Combining \emphCaenorhabditis elegans High-Throughput Fitness Assays with a Large Collection of Recombinant Strains. G3 5, g3.115.017178–920 (2015).

29. Y. V. Makeyeva, M. Shirayama, C. C. Mello, Cues from mRNA splicing prevent default Argonaute silencing in C. elegans. Dev. Cell 56, 2636–2648.e4 (2021).

30. W.-S. Wu, W.-C. Huang, J. S. Brown, D. Zhang, X. Song, H. Chen, S. Tu, Z. Weng, H.-C. Lee, pirScan: a webserver to predict piRNA targeting sites and to avoid transgene silencing in C. elegans. Nucleic Acids Res. 46, W43–W48 (2018).

31. D. Zhang, S. Tu, M. Stubna, W.-S. Wu, W.-C. Huang, Z. Weng, H.-C. Lee, The piRNA targeting rules and the resistance to piRNA silencing in endogenous genes. Science 359, 587–592 (2018).

32. W. Tang, S. Tu, H.-C. Lee, Z. Weng, C. C. Mello, The RNase PARN-1 Trims piRNA 3’ Ends to Promote Transcriptome Surveillance in C. elegans. Cell 164, 974–984 (2016).

33. U. Seroussi, A. Lugowski, L. Wadi, R. X. Lao, A. R. Willis, W. Zhao, A. E. Sundby, A. G. Charlesworth, A. W. Reinke, J. M. Claycomb, A comprehensive survey of C. elegans argonaute proteins reveals organism-wide gene regulatory networks and functions. Elife 12 (2023).

34. W. Gu, M. Shirayama, D. Conte Jr, J. Vasale, P. J. Batista, J. M. Claycomb, J. J. Moresco, E. M. Youngman, J. Keys, M. J. Stoltz, C.-C. G. Chen, D. A. Chaves, S. Duan, K. D. Kasschau, N. Fahlgren, J. R. Yates 3rd, S. Mitani, J. C. Carrington, C. C. Mello, Distinct argonaute-mediated 22G-RNA pathways direct genome surveillance in the C. elegans germline. Mol. Cell 36, 231–244 (2009).

35. F. K. Nelson, D. L. Riddle, Functional study of the Caenorhabditis elegans secretory-excretory system using laser microsurgery. J. Exp. Zool. 231, 45–56 (1984).

36. W. C. Forrester, G. Garriga, Genes necessary for C. elegans cell and growth cone migrations. Development 124, 1831–1843 (1997).

37. S. Liégeois, A. Benedetto, G. Michaux, G. Belliard, M. Labouesse, Genes required for osmoregulation and apical secretion in Caenorhabditis elegans. Genetics 175, 709–724 (2007).

38. M. P. Bagijn, L. D. Goldstein, A. Sapetschnig, E.-M. Weick, S. Bouasker, N. J. Lehrbach, M. J. Simard, E. A. Miska, Function, targets, and evolution of Caenorhabditis elegans piRNAs. Science 337, 574–578 (2012).

39. A. Ashe, A. Sapetschnig, E.-M. Weick, J. Mitchell, M. P. Bagijn, A. C. Cording, A.-L. Doebley, L. D. Goldstein, N. J. Lehrbach, J. Le Pen, G. Pintacuda, A. Sakaguchi, P. Sarkies, S. Ahmed, E. A. Miska, piRNAs can trigger a multigenerational epigenetic memory in the germline of C. elegans. Cell 150, 88–99 (2012).

40. H.-C. Lee, W. Gu, M. Shirayama, E. Youngman, D. Conte Jr, C. C. Mello, C. elegans piRNAs mediate the genome-wide surveillance of germline transcripts. Cell 150, 78–87 (2012).

41. M. Shirayama, M. Seth, H.-C. Lee, W. Gu, T. Ishidate, D. Conte Jr, C. C. Mello, piRNAs initiate an epigenetic memory of nonself RNA in the C. elegans germline. Cell 150, 65–77 (2012).

42. A. E. Sundby, R. I. Molnar, J. M. Claycomb, Connecting the Dots: Linking Caenorhabditis elegans Small RNA Pathways and Germ Granules. Trends Cell Biol. 31, 387–401 (2021).

43. J. Vedanayagam, Small RNA-mediated suppression of sex chromosome meiotic conflicts during Drosophila male gametogenesis. Biochem. Soc. Trans. 53, 281–291 (2025).

44. N. Phadnis, H. A. Orr, A single gene causes both male sterility and segregation distortion in Drosophila hybrids. Science 323, 376–379 (2009).

45. J. Bladen, H.-J. Nam, N. Phadnis, Transformation of meiotic drive into hybrid sterility in Drosophila. Genetics 228, iyae133 (2024).

46. E. C. Andersen, J. S. Bloom, J. P. Gerke, L. Kruglyak, A variant in the neuropeptide receptor npr-1 is a major determinant of \emphCaenorhabditis elegans growth and physiology. 10, e1004156 (2014).

47. E. Ben-David, J. Boocock, L. Guo, S. Zdraljevic, J. S. Bloom, L. Kruglyak, Whole-organism eQTL mapping at cellular resolution with single-cell sequencing. Elife 10 (2021).

48. A. M. Bhagwat, J. Graumann, R. Wiegandt, M. Bentsen, J. Welker, C. Kuenne, J. Preussner, T. Braun, M. Looso, multicrispr: gRNA design for prime editing and parallel targeting of thousands of targets. Life Sci Alliance 3 (2020).

49. R. Core, TEAM, 2017. R: A language and environment for statistical computing. R Foundation for Statistical Computing, Vienna, Austria. Online: https://www.r-project. org (2022).

50. H. Li, Minimap2: pairwise alignment for nucleotide sequences. Bioinformatics 34, 3094–3100 (2018).

51. S. Mao, Y. Qi, H. Zhu, X. Huang, Y. Zou, T. Chi, A Tet/Q Hybrid System for Robust and Versatile Control of Transgene Expression in C. elegans. iScience 11, 224–237 (2019).

52. H.-G. Drost, A. Gabel, I. Grosse, M. Quint, Evidence for active maintenance of phylotranscriptomic hourglass patterns in animal and plant embryogenesis. Mol. Biol. Evol. 32, 1221–1231 (2015).

53. S. Zdraljevic, C. Strand, H. S. Seidel, D. E. Cook, J. G. Doench, E. C. Andersen, Natural variation in a single amino acid substitution underlies physiological responses to topoisomerase II poisons. PLoS Genet. 13, e1006891 (2017).

54. W. A. Boyd, M. V. Smith, J. H. Freedman, Caenorhabditis elegans as a model in developmental toxicology. Methods Mol. Biol. 889, 15–24 (2012).

55. T. C. Shimko, E. C. Andersen, COPASutils: an R package for reading, processing, and visualizing data from COPAS large-particle flow cytometers. PLoS One 9, e111090 (2014).

