## Supplemental Materails for "Divergent *C. elegans* toxin alleles are suppressed by distinct mechanisms"

### List of Supplementary Materials:

Materials and Methods
Fig. S1
Fig. S2
Fig. S3
Fig. S4
Fig. S5
Table S1: Strains
Table S2: Plasmids
Table S3: Plasmid Construction
Table S4: Crosses
Table S5: Oligos
Table S6: gRNAs and repair
Table S7: Inducible construct phenotypes
Table S8: COPAS biosort data

### Materials and methods

#### Strain maintenance

Unless otherwise specified, all strains were propagated at 20 ºC on a modified nematode growth medium (NGMA) containing 1% agar and 0.7% agarose and fed *Escherichia coli* strain OP50 (*46*). Strain names and genotypes can be found in Supplementary Table 1.

#### Plasmids

Plasmid descriptions can be found in Supplementary Table 2. All plasmids generated in this study (with the exception of pWM17) were assembled using gene sequences ordered from IDT (gBlocks) and plasmid backbones (ordered from Addgene or derived from an existing Addgene plasmid) using the NEB Gibson Assembly Master Mix (#E2611). Gibson assemblies were transformed the assembly mix into DH5α competent cells (NEB #C2987H). Plasmid backbones were digested with restriction enzymes and purified using Qiagen QIAquick gel extraction kit (#28706). The gBlocks ordered from IDT, the restriction enzymes used, and the primers used to add homology arms to backbones can be found in Supplementary Table 3. pWM17 was generated using the NEB Q5 site-directed mutagenesis kit (#E0554S) using pWM11 as the template and primers oZ288 & oZ289. All plasmids were purified using the Invitrogen PureLink HQ Mini Plasmid DNA Purification Kit (#K210001) and verified using whole plasmid sequencing with Primordium.

#### Multi-generational cross

QX2538 (*fog-2*(qq212[P17stop]) in XZ1516), QX2539 (*fog-2*(qq212[P17stop]) in QX1211), and QX2327 (qqIr39[fog-2(q71), N2 > DL238] V) were used to generate the cross populations. QX2538 males were used to start the QX2538 x QX2539 cross population and QX2327 males were used to start the QX2327 x QX2538 cross population. Crosses were amplified on 10cm NGMA plates until approximately 20,000 worms were obtained, at which point the population size was maintained. For each generation, worms were washed off the plates using M9, bleach synchronized, and arrested overnight as L1s. The following day, 20,000 L1s were plated across ten 10cm plates at ~2000 worms/plate. The crosses were propagated until the 10th generation of intercrossing. Multiple timepoints were saved for sequencing by freezing a pellet of several thousand worms. For generation ten, DNA libraries were generated using the Illumina Nextera XT DNA Library Preparation Kit (#FC-131-1024) and whole genome sequenced using Illumina NextSeq2000 P1 reagents (20074933). For generation four, allele frequencies were inferred as previously described (*47*).

#### Crosses

Descriptions of the crosses and their phenotypes can be found in the Supplementary Table 4. Workflow for crosses varied slightly depending on the use of fluorescent males. For all crosses, males and L4 hermaphrodites were allowed to mate 24-36 hours before potentially mated hermaphrodites were singled out onto fresh plates. For crosses set up with wildtype males, only progeny from plates with a 50:50 male to hermaphrodite ratio were used for subsequent crosses. For crosses set up with fluorescent males, only progeny with the fluorescent marker were used for subsequent crosses. For self crosses, ~10 hermaphrodite cross progeny were transferred to fresh agar plates and allowed to lay embryos for ~24 hours. For parental backcrosses, ~5 L4 cross progeny were allowed to mate with the designated parent for 24-36 hours before potentially mated hermaphrodites were singled out onto fresh plates and laid embryos for ~24 hours. For all crosses, a defined number of embryos were transferred to fresh plates and allowed to hatch overnight. The following day, the number of arrested larvae were counted.

#### NIL construction

The QX2500 and QX2501 NILs were constructed as previously described (*18*). Briefly, QX2500 was constructed by genotyping F2 progeny of a QX1211 x XZ1516 cross. DNA from F2/F3 progeny was amplified using primer pairs: oZ50-51 and oZ52-53, which enabled us to identify recombinants within a defined genomic region. Recombinant progeny identified with these primer pairs were recovered and backcrossed to QX1211 for 6 generations. We generated QX2501 using Cas9-induced non-homologous recombination to induce strand exchange (*18*). Young adult QX2500/QX1211 heterozygotes were injected with four target gRNAs (gSZ63, gSZ65, gSZ68, and gSZ70), the *dpy-10* gRNA and repair template (see Cas9 injections). We transferred F2 *rol* animals to 96-well plates, allowed them to self, and identified recombinant individuals using oZ66-67 and oZ64-65. We identified and isolated a recombinant that we named QX2501, which contains the QX1211 genotype from V:1-21,536,657, followed by a deletion spanning V:21,536,657-21,547,827, and the XZ1516 genotype from V:21,547,828-22,058,188. All genotyping primers can be found in Supplementary Table 5.

#### Fine-mapping the element

We previously described the construction of the 10 gene CINR-generated NIL (*18*). Briefly, young adult QX1211/QX2501 were injected with gSZ71 and progeny were genotyped with oZ80-82-86 and oZ64-65 to identify individuals that underwent a loss of heterozygosity event.

#### Injection workflow

The following workflow was used for all injections (candidate gene KOs, antidote rescue, and tet-inducible system). The day before injections, L4 animals were transferred to fresh 6 cm NGMA plates and allowed to develop overnight. The following day, young adults were injected and transferred to a fresh 6 cm NGMA plate to recover. The injected animals were singled to fresh 6 cm NGMA plates approximately 16 hours after injections and monitored for the co-injection phenotype.

#### Candidate gene knockouts with CRISPR/Cas9

All guide RNAs were designed to target the XZ1516 genome using the *multicrispr* R package (*18*, *48*, *49*). Guide RNAs were purchased from Synthego as synthetic spacer-scaffold fusions. Guide RNAs were resuspended in 30 µl of water (50 µM) and stored at -20ºC. Cas9 was purchased from IDT (cat #1081059) and stored in single-use aliquots (0.5 µl ~ 5 µg Cas9) at -80ºC. Injection mixtures were made on ice immediately before use. For each mixture, gRNAs (including oZ30; *dpy-10*) were added to a Cas9 aliquot and incubated at 37ºC for 10 minutes. Then the *dpy-10* single-stranded oligodeoxynucleotide (ssODN) repair template (oZ31) and water were added to a final volume of 20 µl. The mixture was spun down in a table-top centrifuge at maximum speed for five minutes and 10µl was taken off of the top to use in injections. The final concentrations in the injection mixtures were: 1.5 µM of Cas9, 4.45 µM gRNA (with each gRNA represented equally), and 0.5 µM of the ssODN repair template. gRNA and repair template sequences can be found in Supplementary Table 6.

#### Antidote rescue

The antidote rescue injection mixture was made on the same day as injections. The mixture was made at room temperature, spun down at maximum speed for five minutes, and 10µl was taken off of the top to use in injections. The final concentrations were: 30 µM pWM4, 5 µM pCFJ104, and 65 µM GeneRuler 1kb DNA ladder (CAT#SM0311). Both DL238 and XZ1516 were injected. Progeny of injected animals that expressed the co-injection marker pCFJ104 were isolated and propagated. DL238 co-inj (+) animals were crossed to XZ1516 WT and XZ1516 co-inj (+) animals were crossed to DL238 WT to assess lethality. In addition to the previously described cross workflow, we also took note of whether or not dead animals inherited the array (as indicated by co-inj (+)).

#### Long read direct RNA sequencing

We extracted and purified RNA (MasterPure #MC85200) from a mixed stage culture of XZ1516. A TapeStation confirmed the high quality of the sample (RINe = 9.3). We prepared long read RNA libraries using Oxford Nanopore library prep kit #SQK-RNA002 on the purified XZ1516 RNA. Libraries were run on a MinION Flow Cell (R9.4.1), bases were called using *guppy*, and the long reads were aligned to the XZ1516 genome using *minimap2 (50)*.

#### Tet-inducible system

We used a tet-inducible system to test the toxicity of several constructs (*51*). The system consists of three plasmids: a tet promoter driving the CDS of interest, a tet activator (TC374), and a tet GFP (TC358). Injection mixtures were made at room temperature, spun down at maximum speed for five minutes, and 10µl was taken off of the top to use in injections. The final concentrations were: 5 µM expression construct (pWM8, 11, 12, 17, or 21), 5 µM TC374, 5µM TC358, 5 µM pCFJ104, and 80 µM GeneRuler 1kb DNA ladder (CAT#SM0311). DL238 was injected with pWM11, pWM12, and pWM17 to test *tmrl-1* toxicity. DL238, XZ1516, and N2 were injected with pWM21 to test *B0250.8* toxicity and with pWM8 as a negative control for all tet-inducible injections. Progeny of injected animals that expressed the co-injection marker pCFJ104 were isolated and propagated. We assessed toxicity of the expression construct of interest through induction on NGM plates with 0.17% doxycycline hyclate (Sigma # D9891), seeded with HT115 bacteria. Most often pCFJ104 (+) gravid worms were bleached onto dox and control plates and phenotyped 48 hours later.

#### Microscopy

RNA FISH: The sequence corresponding to the *tmrl-1* transcript was uploaded to Stellaris's probe design feature, which yielded a mixture of 25 RNA probes, each 19 nucleotides long (Biosearch Technologies). Probes were labeled with CAL Fluor Red 610. XZ1516, DL238, and QX2513 were grown in liquid cultures to amplify the populations. Gravid adults were bleached from liquid cultures and embryos were fixed immediately or allowed to arrest in liquid overnight. Embryos and arrested L1s were prepared according to the Stellaris RNA FISH Protocol for *C. elegans* with the following modifications. Rather than using chambered coverglass, the hybridization and subsequent wash steps were conducted on fixed embryos or larvae in a microcentrifuge tube. ProLong™ Glass Antifade Mountant (ThermoFisher #P36982), rather than Vectashield mounting medium, was added to prepared samples before mounting on slides. Images were taken using a NIKON Eclipse Ti2 widefield microscope with a 100X/1.45 plan apochromat lambda D oil objective, a Photometrics Prime 95B large field of view monochrome fluorescence camera, and a SpectraIII/Celesta/Ziva, MultiLaser(SpectraIII/Lida) light source. The following filters were used: UV excitation 365nm, emission 435nm, and mCherry excitation 514nm, emission 543nm. z-stack images were taken with a 0.2µm step size. Image analysis was performed in NIS Elements AR5.4.

#### Population analysis of the TA element

To determine the strain relatedness of the *C. elegans* population, we extracted variants in the region surrounding the B0250.4 and B0250.8 (V:20454811-20473950) from the CeNDR VCF (version - 20220216). After subsetting the VCF, we used the *vcf2dist* and *dist2tree* functions in the *fastreeR* R package to generate the relatedness dendrogram.

We used the R package *orthologr* to calculate nucleotide and amino acid identity and dS (*52*). We calculated dS across all orthologs between the N2 and XZ1516 genomes using all of the methods available in *orthologr*. We used the following formula to calculate divergence time: $T = (dS - \pi anc)/(2\mu)$ (*21*), as previously described (*22*).

#### Mut-16 phenotyping

QX2537, QX2532, and QX5434 were propagated on 10cm NGMA plates until gravid. The three strains were bleached and L1s were arrested overnight shaking at 20ºC and 180 rpm in K media (*53*, *54*). The next day, samples were fed HB101 E. coli at a final concentration of OD10 and allowed to grow for 48 hours, shaking at 20ºC and 180 rpm. Prior to scoring strains on the COPAS BIOSORT using the sample cup, sodium azide was added at a final concentration of 50 mM (*28*). The resulting data was loaded into R for analysis (*49*). Objects with a TOF value greater than 60 and less than 1000 were retained for analysis. We used a pre-trained SVM to detect bubbles in the data set (*COPASutils::bubbleSVMmodel_noProfiler*; threshold = 0.9999999) (*55*). Larval stages were defined by TOF (60 < TOF < 90 = L1; 90 < TOF < 200 = L2/L3; 200 < TOF < 300 = L4; 300 < TOF < 1000 = adult). We note that the conclusions drawn from this experiment did not depend on the SVM threshold used (Fig. S6)

### Supplementary Figures:

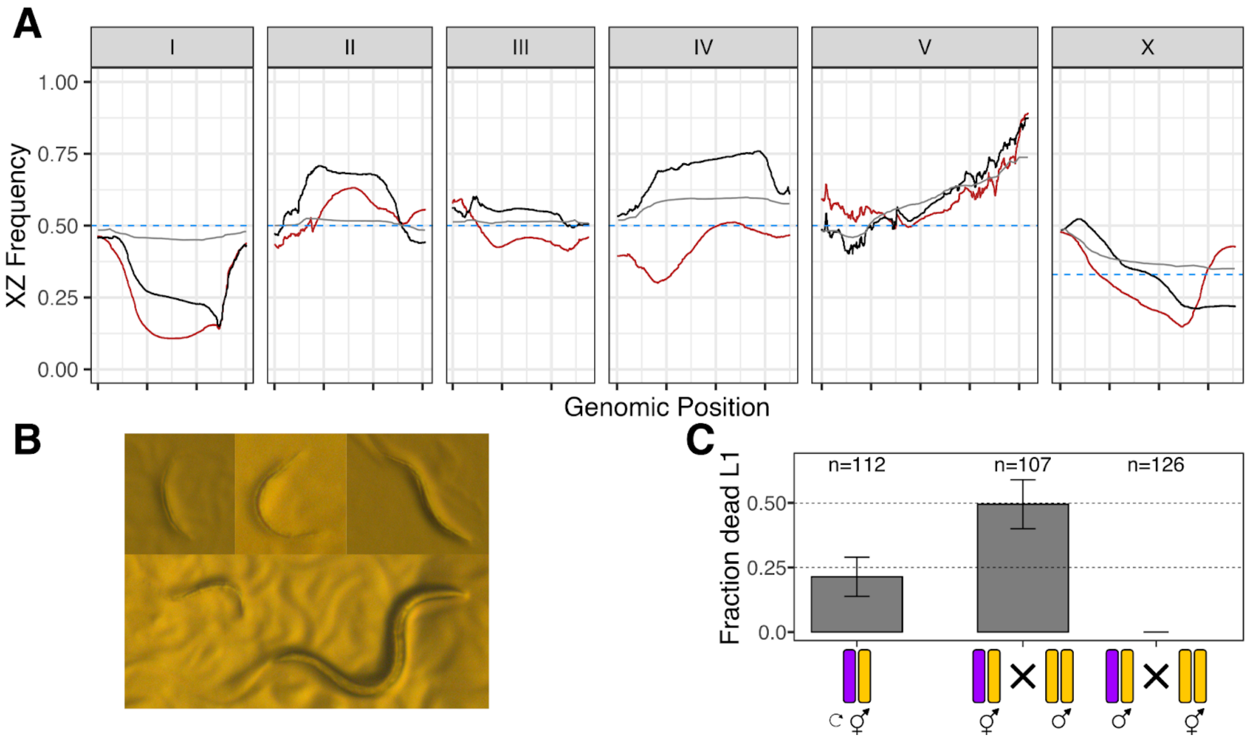

**Fig. S1. DL238 is also susceptible to the XZ1516 TA**

A) Estimates of the XZ1516 allele frequency at each position in the genome after four (gray) or ten generations of intercrossing with QX1211 (black, gray) or DL238 (red). Each panel corresponds to a *C. elegans* chromosome and each x-axis tick represents 5 Mb. The dotted blue line represents the expected allele frequency for each chromosome. B) Top three panels are representative images of *rod*-like lethality phenotype caused by the *tmrl-1* toxin. The bottom panel is a *rod* worm next to a normally developing worm derived from the same cross. C) Crosses between XZ1516 (purple) and DL238 (yellow) that establish the inheritance pattern of the TA element. The y-axis represents the fraction of dead L1s observed for each cross depicted on the x-axis. Crosses from left to right: selfing of XZ1516/DL238 heterozygous; XZ1516/DL238 heterozygous hermaphrodites crossed to DL238 males; XZ1516/DL238 heterozygous males crossed to DL238 hermaphrodites. The observed fraction of dead L1s was not significantly different from the expected fractions for a maternally inherited TA element, exact binomial test.

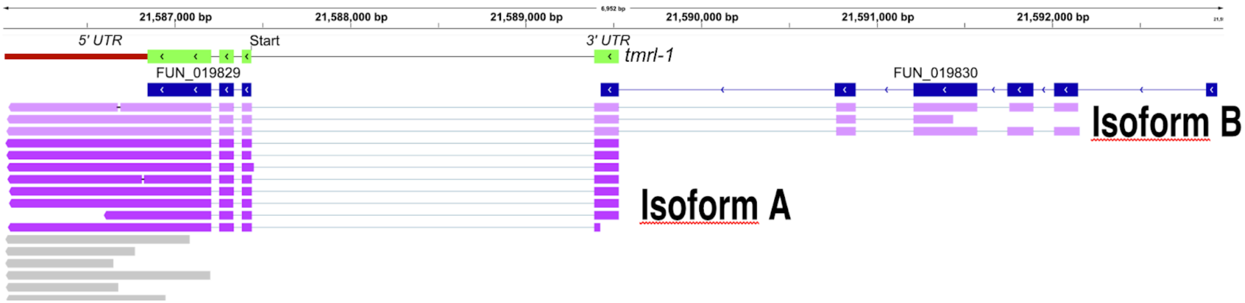

**Fig. S2. Isoforms of the *tmrl-1* toxin**
Long-read RNA sequencing reads of the different *tmrl-1* isoforms aligned to the XZ1516 genome visualized in the Integrative Genomics Viewer (IGV). Light purple reads represent isoform B, the dark purple reads represent isoform A, and gray reads represent the 5'UTR of *tmrl-1*. The functional gene model of *tmrl-1* is depicted in green above the computationally predicted "FUN" gene models.

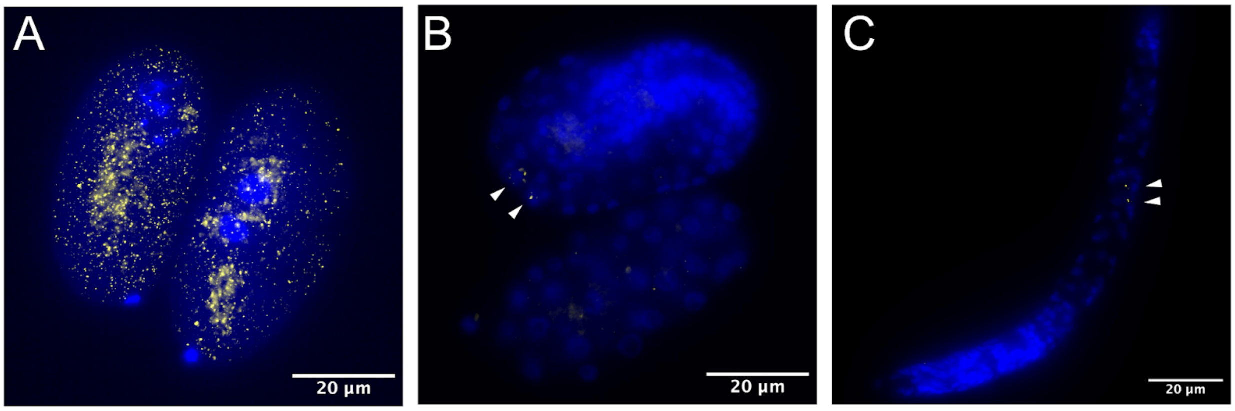

**Fig. S3. The tmrl-1 transcript persists until the L1 stage**

Representative *tmrl-1* FISH images in XZ1516. Yellow corresponds to *tmrl-1* transcript puncta and nuclei staining with DAPI is in blue. A) Max projection of 1- and 2-cell embryos. B) Post-gastrulation embryos showing *tmrl-1* localization to the Z2/Z3 cells. C) L1 larvae showing *tmrl-1* localization to the Z2/Z3 cells.

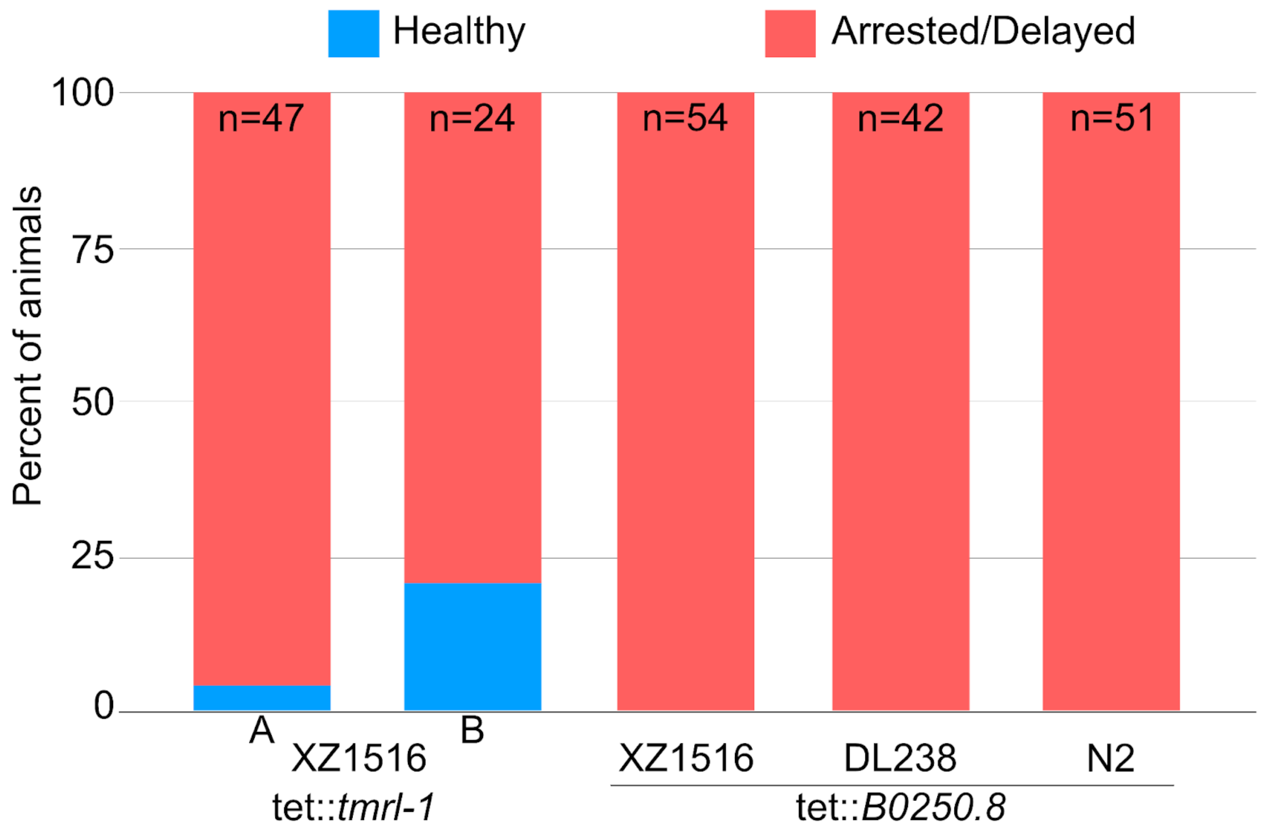

**Fig. S4. Phenotypes induced by the two *tmrl-1* alleles**

The percent of healthy (blue) or arrested or delayed (red) individuals is shown on the y-axis. The first two bars represent technical replicates of tetracycline-induced *tmrl-1* in the XZ1516 genetic background. The remaining three bars correspond to tetracycline-induced *B0250.8* in XZ1516, DL238, and N2, respectively.

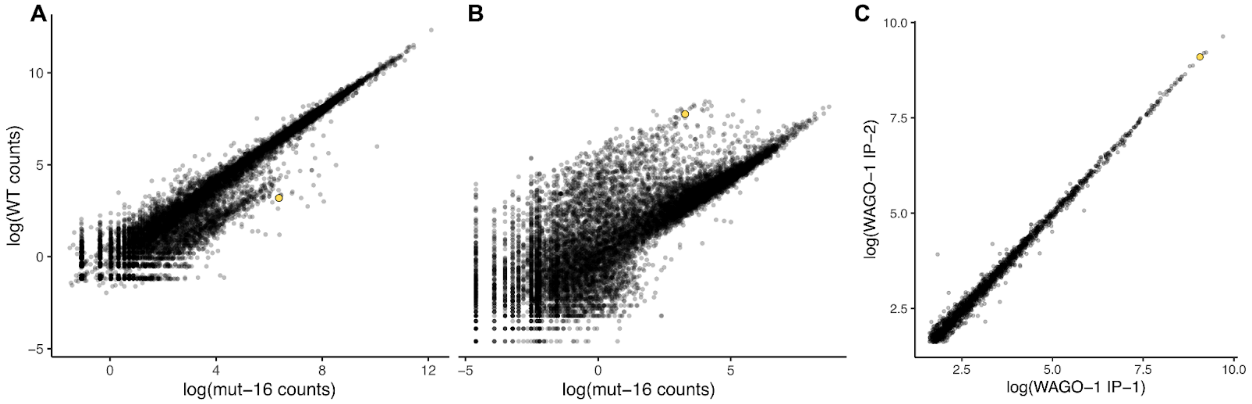

**Fig. S5. Small RNAs are involved in the suppression of the N2 tmrl-1 allele**

The A) mRNA and B) sRNA abundance is shown for Δ*mut-16* on the x axis and WT on the y axis (*26*). The N2 *tmrl-1* (*B0250.8*) allele is represented as a yellow dot. C) sRNA abundance of WAGO-1-associated sRNAs for two independent replicates (*33*).

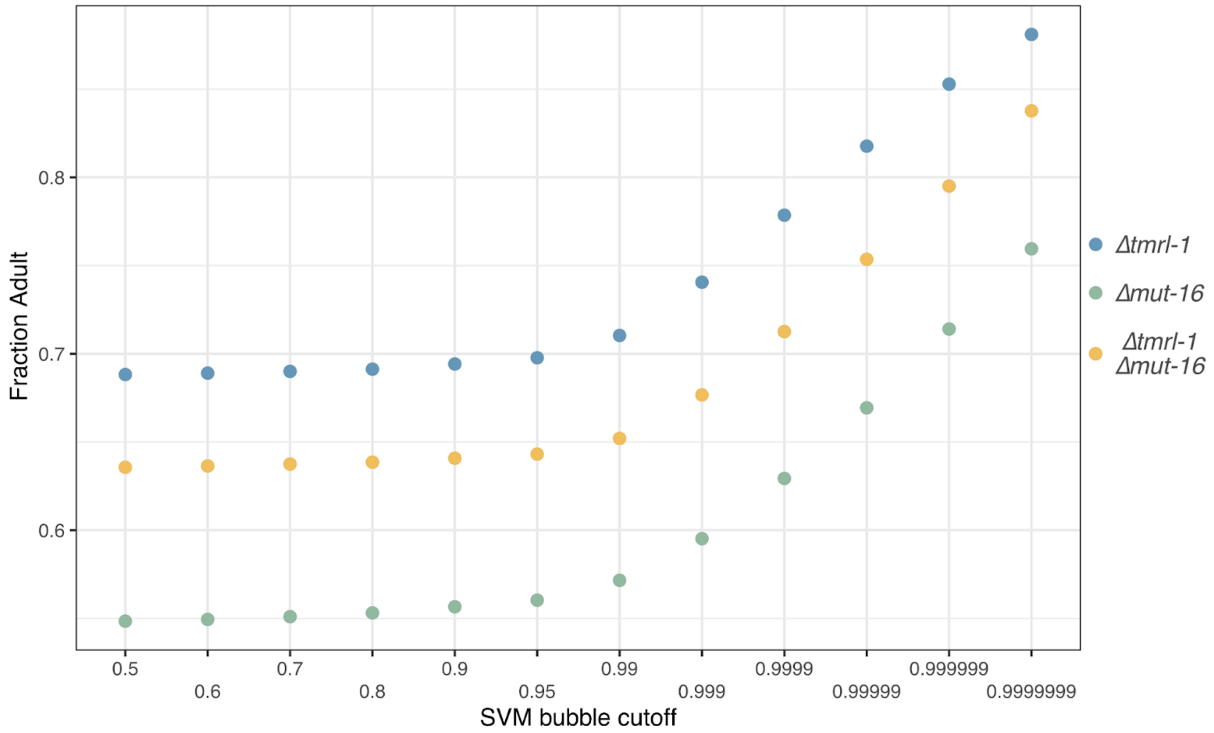

**Fig. S6. SVM bubble detection for COPAS biosort**
The fraction of individual worms that developed to adults is shown on the y axis for increasingly strict SVM cutoffs that differentiate worms from bubbles on the x axis. The most strict cutoff of 0.9999999 was used to generate Fig. 4.

### Supplementary Tables:

#### Table S1

| Strain | Allele name | Genotype | Description |
|---|---|---|---|
| XZ1516 | | | Wild C. elegans strain |
| QX1211 | | | Wild C. elegans strain |
| DL238 | | | Wild C. elegans strain |
| N2 | | | C. elegans reference strain |
| QX2538 | qq212 | *fog-2(qq212[P17stop])* in XZ1516 | XZ1516 with null *fog-2*; used to generate large cross population with QX1211 |
| QX2539 | qq212 | *fog-2(qq212[P17stop])* in QX1211 | QX1211 with null *fog-2*; used to generate large cross population with XZ1516 |
| QX2327 | qqIR39 | qqIr39[fog-2(q71), N2 > DL238] V | qqIr38 is an introgression of the *fog-2*(q71) allele into DL238 |
| QX2500 | qqIR50 | qqIR50(V:18,699,989-22,058,188, XZ1516>QX1211) | NIL used to generate QX2501 recombinant |
| QX2501 | qqIR51 | qqIR51(V:21,547,828-22,058,188, XZ1516>QX1211) | NIL used to generate CINR recombinants. Contains a deletion spanning V:21537348-21547827 |
| QX2506 | qqIR56 | qqIR56(V:21,547,828-21,594,790, XZ1516>QX1211) | CINR recombinant generated with gSZ71 (XQ) |
| QX2509 | qq200 | FUN_019827-FUN_019830(qq200) in XZ1516 | Cas9 induced deletion (gSZ85 & gSZ86) in XZ1516 V:21,583,509-21,591,326; partial deletion FUN_019827 & FUN_019830 and full deletion FUN_019828 & *tmrl-1* |
| QX2510 | qq201 | FUN_019827-FUN_019830(qq201) in XZ1516 | Cas9 induced deletion (gSZ85 & gSZ86) spanning XZ1516 genome coordinates V:21,583,481-21,591,295; partial deletion FUN_019827 & FUN_019830 and full deletion FUN_019828 & *tmrl-1* |
| QX2511 | qq202 | *tmrl-1*(qq202) in XZ1516 | Cas9 induced deletion (gSZ83 & gSZ108) spanning XZ1516 genome coordinates V:21,586,947-21,587,463; full deletion *tmrl-1* |
| QX2512 | qq203 | *tmrl-1*(qq203) in XZ1516 | Cas9 induced deletion (gSZ83 & gSZ108) spanning XZ1516 genome coordinates V:21,586,947-21,587,434; full deletion *tmrl-1* |
| QX2513 | qq204 | *tmrl-1*-FUN_019830(qq204) in XZ1516 | Cas9 induced deletion (gSZ83, gSZ85 & gSZ87) spanning XZ1516 genome coordinates |

| | | | |
|---|---|---|---|
| | | | V:21,586,948-21,592,275; full deletion *tmrl-1* & FUN_019830 |
| QX2514 | qq205 | *tmrl-1*-FUN_019830(qq205) in XZ1516 | Cas9 induced deletion (gSZ83, gSZ85 & gSZ87) spanning XZ1516 genome coordinates V:21,586,952-21,592,809; full deletion *tmrl-1* & FUN_019830 |
| QX2515 | qq206 and qq207 | qq206 [KI eft-3p::mCherry::eft-3utr, IV:18146714], qq207 [KI eft-3p::GFP::eft-3utr HygR, IV:13050294] in XZ1516 | Fluorescent XZ1516 used to verify cross progeny; COP2548 generated by InVivo Biosystems backcrossed to XZ1516 6x |
| QX2516 | qq203 and qq208 | *tmrl-1*(qq203), *amrl-1*(qq208) in XZ1516 | Cas9 induced deletion [V:21,565,726-21,573,347] using gSZ111 & gSZ112 in strain QX2512; full deletion *tmrl-1* & *amrl-1* |
| QX2517 | qq203 and qq209 | *tmrl-1*(qq203), *amrl-1*(qq209) in XZ1516 | Cas9 induced deletion [V:21,565,726-21,573,345] using gSZ111 & gSZ112 in strain QX2512; full deletion *tmrl-1* & *amrl-1* |
| QX2518 | qqEx100 | qqEx100[*eef-1A.1p::amrl-1::tbb-2; myo-3p::mCherry::unc-54*] in XZ1516 | XZ1516 injected with pWM4 (to test constitutive expression of *amrl-1*) and pCFJ104 (co-injection marker) |
| QX2519 | qqEx100 | qqEx100[*eef-1A.1p::amrl-1::tbb-2; myo-3p::mCherry::unc-54*] in XZ1516 | XZ1516 injected with pWM4 (to test constitutive expression of *amrl-1*) and pCFJ104 (co-injection marker) |
| QX2520 | qqEx100 | qqEx100[*eef-1A.1p::amrl-1::tbb-2; myo-3p::mCherry::unc-54*] in DL238 | DL238 injected with pWM4 (to test constitutive expression of *amrl-1*) and pCFJ104 (co-injection marker) |
| QX2521 | qqEx100 | qqEx100[*eef-1A.1p::amrl-1::tbb-2; myo-3p::mCherry::unc-54*] in DL238 | DL238 injected with pWM4 (to test constitutive expression of *amrl-1*) and pCFJ104 (co-injection marker) |
| QX2524 | qqEx102 | qqEx102[*TRE3GV::tmrl-1.a; TRE::GFP; Prpl-28::rtetR-QFAD::P2A::mKate::T2A::tetR-pie1*] in XZ1516 | XZ1516 injected with pWM11, TC358, and TC374 (tet inducible system to test the toxicity of *tmrl-1* isoform A) and pCFJ104 (co-injection marker) |
| QX2525 | qqEx103 | qqEx103[*TRE3GV::tmrl-1.a[G43stop]; TRE::GFP; Prpl-28::rtetR-QFAD::P2A::mKate::T2A::tetR-pie1*] in DL238 | DL238 injected with pWM17, TC358, and TC374 (tet inducible system to test the toxicity of *tmrl-1* isoform A with a stop codon at position 43) and pCFJ104 (co-injection marker) |
| QX2526 | qqEx104 | qqEx104[*TRE3GV::tmrl-1.b; TRE::GFP; Prpl-28::rtetR-QFAD::P2A::mKate::T2A::tetR-pie1*] in DL238 | DL238 injected with pWM12, TC358, and TC374 (tet inducible system to test the toxicity of *tmrl-1* isoform B) and pCFJ104 (co-injection marker) |

| | | | |
|---|---|---|---|
| QX2527 | qqEx105 | qqEx105[*TRE3GV::B0250.8; TRE::GFP; Prpl-28::rtetR-QFAD::P2A::mKate::T2A::tetR-pie1*] in XZ1516 | XZ1516 injected with pWM21, TC358, and TC374 (tet inducible system to test the toxicity of *B0250.8*) and pCFJ104 (co-injection marker) |
| QX2528 | qqEx105 | qqEx105[*TRE3GV::B0250.8; TRE::GFP; Prpl-28::rtetR-QFAD::P2A::mKate::T2A::tetR-pie1*] in DL238 | DL238 injected with pWM21, TC358, and TC374 (tet inducible system to test the toxicity of *B0250.8*) and pCFJ104 (co-injection marker) |
| QX2529 | qqEx105 | qqEx105[*TRE3GV::B0250.8; TRE::GFP; Prpl-28::rtetR-QFAD::P2A::mKate::T2A::tetR-pie1*] in N2 | N2 injected with pWM21, TC358, and TC374 (tet inducible system to test the toxicity of *B0250.8*) and pCFJ104 (co-injection marker) |
| QX2530 | qq210 | *amrl-1*(qq210) in NIC195 | Cas9 induced deletion (gSZ78 & gSZ128) spanning XZ1516 genome coordinates V: 21,567,654-21,570,740; large deletion of amrl-1 in NIC195 |
| QX2537 | qq211 | *B0250.8*(qq211) in N2 | Cas9 induced deletion [V:20,469,883-20,470,535] using gSZ119 & gSZ122 in strain N2; full deletion of *B0250.8* |
| QX2532 | qq211 and cmp185 | *B0250.8*(qq211) and *mut-16*(cmp185) in N2 | USC1148 crossed to QX2537 and selected F2 progeny homozygous for qq211 allele and cmp185 allele |
| QX2534 | cmp185 | *mut-16(cmp185[mut-16ΔE-K::mCherry::2xHA]) I.* | USC1148 crossed to QX2537 and selected F2 progeny homozygous for qq211 allele |

#### Table S2

| Plasmid name | Genotype | Description | Source |
|---|---|---|---|
| **pCFJ104** | Pmyo-3::mCherry::unc-54 | Used as a co-injection marker for tet-inducible system | pCFJ104 was a gift from Erik Jorgensen (Addgene plasmid # 19328 ; http://n2t.net/addgene:19328 ; RRID:Addgene_19328) |
| **pWM1** | Peef-1A.1::blank::tbb-2 | Used to build pWM3 and pWM4 | derived from Addgene plasmid # 46168 |
| **pWM4** | Peef-1A.1::amrl-1::tbb-2 | Used to test constitutive expression in the amrl-1 rescue experiment | this study |
| **TC374** | Prpl-28::rtetR-QFAD::P2A::mKate::T2A::tetR-pie1 | Activator construct in the tet-inducible system | TC374 was a gift from Tian Chi (Addgene plasmid # 164886 ; http://n2t.net/addgene:164886 ; RRID:Addgene_164886) |
| **TC358** | TRE::GFP | GFP construct in the tet-inducible system | TC358 was a gift from Tian Chi (Addgene plasmid # 164885 ; http://n2t.net/addgene:164885 ; RRID:Addgene_164885) |
| **pWM8** | TRE::blank | Used to build expression constructs in the tet-inducible system (pWM11,12,17,21,25). Used in the control injection for the tet-inducible system | derived from TC358 |
| **pWM11** | TRE3GV::tmrl-1.a | tet-inducible expression of tmrl-1 isoform a | this study |
| **pWM12** | TRE3GV::tmrl-1.b | tet-inducible expression of tmrl-1 isoform b | this study |
| **pWM17** | TRE3GV::tmrl-1.a[G43stop] | tet-inducible expression of tmrl-1 isoform a with a stop codon introduced at amino acid position 43 | this study |
| **pWM21** | TRE3GV::B0250.8 | tet-inducible expression of B0250.8 | this study |
| **pWM25** | TRE3GV::amrl-1 | tet-inducible expression of amrl-1 | this study |

### Table S3

| Plasmid | Backbone | Insert name | Insert sequence | Insert amplified with | Other |
|---|---|---|---|---|---|
| **pWM4** | pWM1(BsrGI&BamHI) | amrl-1 | ATGACGTCGTTTATGATAAACAAGAAGCAACTGAAAGA<br>AAAGATCCGGCAGCTTCAGGATCGGCTCGAAAGTGGA<br>GTATTCAATAAACAAACTGGAAACAATGGAAACCAGAA<br>AATGCAATTTCAATGCAGTCAGTGTTCAGGTATCTTTCG<br>AAATGAAGACCATCTATGGACTCATAACACGGAATGTCA<br>CAGATCGAGGGATGCCCAAAGAATACCGGAATACTTTT<br>GTGATCAATGCCCAGTGCGGTATGATTACAAATTTCAGT<br>TGATTCGTCACCTGGATTTCGGCCATGAAGAGATAAAG<br>TTCCCGCGGAAACAAGCGACTACAAACAAAGAGGAGT<br>TGTACATGAAACATTACATTAGTGCAAGCGATTTCTATG<br>CAAAAAATCCCGACGCATGTGACAGATGTTACTGGTTC<br>ATTGGAACACCGGAAAAACTGTTGGAACATAAGAATAA<br>GTATCACCCAGGATGGGAAATTCCTTACGAGGCTCCAG<br>TTTATGATAAAAAAGAGTACGTCTACAAGTGTATTTATTG<br>CACAGGAGCATTCCGTTTGGAAGAAGACTTGGTTTACC<br>ATCAAGAAGATTGGCATGAAAACAAAGAAAAACTGAGA<br>AGACATAGTGCTTTACTCCGCCCAACTCACTGTGACAC<br>TTCAGATTGTGCTAAAAGATTCGAGACAAGAGAACAATT<br>GTGGGAACACAGATCGCTTGATCATTCTGACTTTTACG<br>GAAAGCTGGAATGTTTAAATAAAAAAGAATACAAAGATT<br>TTAAAGAAGAAGAAATCCACGGAACCATTGAAGAACTT<br>CAGAAAATGTTAAATAATGGACTATATACGGACGAATTTG<br>ATTTTGACATAACGCCATTCATAACTGTTGATGATGGCTT<br>CAAAGATGTGGCAAGACTTCGTCACAAATGCAATCACT<br>GCTCCGGAGCCTTCCAACAGGAATGTATCATGATGTAT<br>CATCTTCAAATATGCCACCcggggaaaaaaaTGGAATTGAG<br>ACCCACCTACTTTTGTGATCGTTGTCCAATGCGGTTTG<br>GAACCGAGTCGGCGTTGAATTTTCACCAATATGAGACA<br>TATGAAGAGGGACTTGACAAGAGAACTGGCAAACGAT<br>CGTGGTATGGGGTGATTGAAGTATCTAAACAACGTAGG<br>GAGCATTACATTTCCGAAGAAGATTTCTACACTAAATTC<br>CCGGGCGCATGTAACAAATGCTTCAGCAATTTTGATAC<br>CGCGGAGCTCATGTTGGAGCACAAAAGCGAGAAGCAT<br>CCAGGATGGGAACACGAGGAGAAACCAAAAGTTTATG<br>CTGACGATGAATATACCTTCTATTGCCAAGAGAAAATAT<br>GCTATGGAGCATTTCTTTTGGAGCAAGACCTTATTTATC<br>ATCAACAAAAAGTTCACCCCAAGCAGCTCAAAACAGA<br>AAATGTAATTTTCTGCAATAAATGTAGTTATGAATATTTCC<br>ACGAAATGAGCGAGTTTTTGGCTCATAGCCACCAACG<br>GCACGACGGAACAGCAACGGGAGTTCCGAGAATGATT<br>AAGCACATCCCGTCAGGTGGCACATCTGTCCTACCAT<br>GTGAAAAAGAAAGCGATGTGAAAACTGGACATGCGGA<br>GAAAGACGACGACACTGACTCCTCTTTTGAGATTGTCT<br>AA | oZ164 & oZ165 | |
| **pWM11** | pWM8(EcoRI) | tmrl-1.a | AGAGGTGCTTCGGACGTCGCATCCGCATCCAGCGTAA<br>AGGAGCTGGTGGAATCTTCAAGAGCCACAACAAGCAC<br>CGCAAAGGAGCATCCAAGCTCCGAACAATAGCCATCG<br>TCGCTCGACATTTCAAGGATATCAAAAAGCCGACATGC | oZ203 & oZ204 | |

| | | | | | |
|---|---|---|---|---|---|
| | | | CTTCTCAGGAAGCCGTTCAAGCACTTGTAGAGACGTTTCGCACAACTCCGGAGAATCCTCTGCTCATCAAATCAGGCATGGAGATGCTCAGTTTCTCATATTCGGAAAAGGAATCGCTAAATGGACACGAGatCAAAATTGACAGCTCCAACGGACGTATCACATTCACCGTTTCGAAAATTGACCACCCGTCCAAATTTGATATTGAGGAGTACATTGGAGTTGTGCAGGAAAACTGCCAAAATGAACCCCATGTTCCTATGCAGCCATCTCCGAATCGCCGCCATGGTGTTGACAAAGGCACAGGAGAAGAGGCGAAGGAAGTCTCCAATGCCACATATTTTGCTAGAATTGTCGATATTTCGGGAAAACCAGACCACTTTGGATCCCGCGGCACATATCTAAGTCCATCGGAGATTGAGATTGAAAAGATCGGATTGAAGAATGATGTAGTCTACGGTCTATATGTCGCTACCAAAACCAGTCCACTCGAATAA | | |
| **pWM12** | pWM8(EcoRI) | tmrl-1.b | TTTCAGGTGATGCACGGGGTTGGCTGTACAACCGTGCAGAGGAGATTGGCACCTCAGGAATGGAAGGAATGGGTCCCTTCCATTCCCGCCTCTCCGGTGTTGCAGGTGAAATTGTTTGCCTGGGCACCGGAAAAGAAACTTGGGTAACACGGACTTCGTGTCCAGTGCAAAGGAGCCGGTGGAATCTTCAAGAGCCACAACAAGCACCGCAAAGGAGCATCCAAGCTCCGTCCTCTCGACTATGCCGAGAGACATGGCTACATCAAAGGATTCGTGaattaggttaataaGGGTGAACGTCTCCTGCACGCTGTCGGCTGTACATTATCAATTCTTTGAGAGAAAGAAAAGACGAGACGAAGAAACAGCTCAAGTAAATGTGCAATCAGCTCAAGAATCAACTTCATCCACTCCACCGAAAGACATATTCCACCCAAATTGGGAGGATACAACATCAGAAGGAACTTCAACTCAAAACCCTTCGAATTCCCGTTCAAACCTGGCTTATCAGTCGTTCCGGAAGCACGTTTCAATTGAAGATCGATATTTGGATCCACAAGCAGTAATTATTAACGAGGTGCCGGCATCAAATCATATGGATGAGATTAGAAGGATATCGAAGCCGAGACCCACTGGAAATCCGGCTCTGAGACAGGGTTCCCTTCGAGCACCACCTGATATGCCATCGTCTTCTGATCAATTCAAGTGAAATATGTCGTTAAGGATCGTGTCGAATAATCCAACGGTAGATCAGACGACTGAAACGAGGTGCTTCGGACGTCGCATCCGCATCCAGCGTAAAGGAGCTGGTGGAATCTTCAAGAGCCACAACAAGCACCGCAAAGGAGCATCCAAGCTCCGAACAATAGCCATCGTCGCTCGACATTTCAAGGATATCAAAAAGCCGACATGCCTTCTCAGGAAGCCGTTCAAGCACTTGTAGAGACGTTTCGCACAACTCCGGAGAATCCTCTGCTCATCAAATCAGGCATGGAGATGCTCAGTTTCTCATATTCGGAAAAGGAATCGCTAAATGGACACGAGatCAAAATTGACAGCTCCAACGGACGTATCACATTCACCGTTTCGAAAATTGACCACCCGTCCAAATTTGATATTGAGGAGTACATTGGAGTTGTGCAGGAAAACTGCCAAAATGAACCCCATGTTCCTATGCAGCCATCTCCGAATCGCCGCCATGGTGTTGACAAAGGCACAGGAGAAGAGGCGAAGGAAGTCTCCAATGCCACATATTTTGCTAGAATTGTCGATATTTCGGGAAAACCAGACCACTTTGGATCCCGCGGCACATATCTAAGTCCATCGGAGATTGAGATTGAAAAGATCGGATTGAAGAATGATGTAGTCTACGGTCTATATGTCGCTACCAAAACCAGTCCACTCGAATAA | oZ205 & oZ204 | |
| **pWM17** | NA | NA | NA | NA | Q5 site-directe |

| | | | | | |
|---|---|---|---|---|---|
| | | | | | d mutagenesis: pWM11 amplified with oZ288 & oZ289 |
| **pWM21** | pWM8(EcoRI) | B0250.8 | ATGCCAGCACAAAAAGGAGTACAAGCCCTTGTATCAAAGTTTCGTCTTTCTCCAGGTAACCCAATGTTCCTGAAATCTAGTTCAGATTTATTCAGCTTCACTTCTTCAGACAAAGAATATCTGGAAGGTTATGAGATAAAGGTCAAAAGCGCCGCAGGAGTTATCAAATTCACTCTCTTGAAAGGTGATCAACCGCCTAAATATGAAATCAGTTCTTTCCTTGGAATCATTTTGGAACACCGTGAAGATCCAAGTAATACGCCGATGCCGCCGTCTTCGAGTCGCCGTCATGCAGTGGACAATGGCAAGGGAGAAGAAGCGAAGTCTGTCACCAATGACACAGTTTTCGCACGAATTCAGGATTACTTTGGAAAACAAATGCACGTTGGCAGTTATGGCTCAACATTCAAACCATCGAAGATTGAGATAAACGAGACCAATGTGAAGAAGGGTCAAGTCTACGGTATTTACCTCGTTTCCAACACCACTTCTTCTGTATAA | oZ217 & oZ218 | |
| **pWM25** | pWM8(EcoRI) | amrl-1 | ATGACGTCGTTTATGATAAACAAGAAGCAACTGAAAGAAAAGATCCGGCAGCTTCAGGATCGGCTCGAAAGTGGAGTATTCAATAAACAAACTGGAAACAATGGAAACCAGAAAATGCAATTTCAATGCAGTCAGTGTTCAGGTATCTTTCGAAATGAAGACCATCTATGGACTCATAACACGGAATGTCACAGATCGAGGGATGCCCAAAGAATACCGGAATACTTTTGTGATCAATGCCCAGTGCGGTATGATTACAAATTTCAGTTGATTCGTCACCTGGATTTCGGCCATGAAGAGATAAAGTTCCCGCGGAAACAAGCGACTACAAACAAAGAGGAGTTGTACATGAAACATTACATTAGTGCAAGCGATTTCTATGCAAAAAATCCCGACGCATGTGACAGATGTTACTGGTTCATTGGAACACCGGAAAAACTGTTGGAACATAAGAATAAGTATCACCCAGGATGGGAAATTCCTTACGAGGCTCCAGTTTATGATAAAAAAGAGTACGTCTACAAGTGTATTTATTGCACAGGAGCATTCCGTTTGGAAGAAGACTTGGTTTACCATCAAGAAGATTGGCATGAAAACAAAGAAAAACTGAGAAGACATAGTGCTTTACTCCGCCCAACTCACTGTGACACTTCAGATTGTGCTAAAAGATTCGAGACAAGAGAACAATTGTGGGAACACAGATCGCTTGATCATTCTGACTTTTACGGAAAGCTGGAATGTTTAAATAAAAAAGAATACAAAGATTTTAAAGAAGAAGAAATCCACGGAACCATTGAAGAACTTCAGAAAATGTTAAATAATGGACTATATACGGACGAATTTGATTTTGACATAACGCCATTCATAACTGTTGATGATGGCTTCAAAGATGTGGCAAGACTTCGTCACAAATGCAATCACTGCTCCGGAGCCTTCCAACAGGAATGTATCATGATGTATCATCTTCAAATATGCCACCcggggaaaaaaTGGAATTGAGACCCACCTACTTTTGTGATCGTTGTCCAATGCGGTTTGGAACCGAGTCGGCGTTGAATTTTCACCAATATGAGACATATGAAGAGGGACTTGACAAGAGAACTGGCAAACGATCGTGGTATGGGGTGATTGAAGTATCTAAACAACGTAGGGAGCATTACATTTCCGAAGAAGATTTCTACACTAAATTCCCGGGCGCATGTAACAAATGCTTCAGCAATTTTGATACCGCGGAGCTCATGTTGGAGCACAAAAGCGAGAAGCATCCAGGATGGGAACACGAGGAGAAACCAAAAGTTTATG | oZ224 & oZ225 | |

| | | | |
|---|---|---|---|
| | | | CTGACGATGAATATACCTTCTATTGCCAAGAGAAAATAT<br>GCTATGGAGCATTTCTTTTGGAGCAAGACCTTATTTATC<br>ATCAACAAAAAGTTCACCCCAAGCAGCTCAAAACAGA<br>AAATGTAATTTTCTGCAATAAATGTAGTTATGAATATTTCC<br>ACGAAATGAGCGAGTTTTTGGCTCATAGCCACCAACG<br>GCACGACGGAACAGCAACGGGAGTTCCGAGAATGATT<br>AAGCACATCCCGTCAGGTGGCACATCTGTCCTACCAT<br>GTGAAAAAGAAAGCGATGTGAAAACTGGACATGCGGA<br>GAAAGACGACGACACTGACTCCTCTTTTGAGATTGTCT<br>AA |

### Table S4

| Cross | Description | Embryos | Alive | Dead L1s | Arrested embryos | Sick worms | Corresponding Figure |
|---|---|---|---|---|---|---|---|
| **QX2515-QX1211_self** | XZ1516(fluorescent males)/QX1211 self | 155 | 112 | 43 | 0 | NA | Fig. 1 |
| **QX2515-QX1211_QX1211 herm** | XZ1516(fluorescent males)/QX1211 male backcrossed to QX1211 hermaphrodite | 104 | 98 | 0 | 6 | NA | Fig. 1 |
| **QX2515-QX1211_QX1211 male** | XZ1516(fluorescent males)/QX1211 backcrossed to QX1211 male | 174 | 93 | 79 | 2 | NA | Fig. 1 |
| **QX2515-DL238_self** | XZ1516(fluorescent males)/DL238 self | 112 | 88 | 24 | 0 | NA | Fig. S1 |
| **QX2515-DL238_DL238herm** | XZ1516(fluorescent males)/DL238 males backcrossed to DL238 hermaphrodites | 126 | 125 | 0 | 1 | NA | Fig. S1 |
| **QX2515-DL238_DL238male** | XZ1516(fluorescent males)/DL238 hermaphrodites backcrossed to DL238 males | 107 | 51 | 53 | 3 | NA | Fig. S1 |
| **QX2500-QX1211_self** | NIL/QX1211 self | 75 | 54 | 21 | 0 | NA | Fig. 2A |
| **QX2500-QX1211_self** | NIL/QX1211 self | 75 | 57 | 18 | 0 | NA | Fig. 2A |
| **QX2500-QX1211_self** | NIL/QX1211 self | 82 | 60 | 22 | 0 | NA | Fig. 2A |
| **QX2501-QX1211_self** | subNIL/QX1211 self | 35 | 28 | 7 | NA | NA | Fig. 2A |
| **QX2501-QX1211_self** | subNIL/QX1211 self | 32 | 22 | 10 | NA | NA | Fig. 2A |
| **QX2501-QX1211_self** | subNIL/QX1211 self | 24 | 16 | 8 | NA | NA | Fig. 2A |
| **QX2501-QX1211_self** | subNIL/QX1211 self | 40 | 34 | 6 | NA | NA | Fig. 2A |
| **QX2506-QX1211_self** | 10-gene NIL/QX1211 self | 36 | 17 | 8 | 11 | NA | Fig. 2A |
| **QX2506-QX1211_self** | 10-gene NIL/QX1211 self | 140 | 110 | 19 | 11 | NA | Fig. 2A |
| **QX2506-QX1211_self** | 10-gene NIL/QX1211 self | 66 | 48 | 15 | 3 | NA | Fig. 2A |
| **QX2506-QX1211_self** | 10-gene NIL/QX1211 self | 77 | 53 | 13 | 11 | NA | Fig. 2A |
| **XZ1516-QX1211_self** | XZ1516/QX1211 self | 47 | 34 | 13 | 0 | NA | Fig. 2B |

| | | | | | | | |
|---|---|---|---|---|---|---|---|
| **XZ1516-QX1211_self** | XZ1516/QX1211 self | 62 | 49 | 13 | 0 | NA | Fig. 2B |
| **XZ1516-QX1211_self** | XZ1516/QX1211 self | 120 | 85 | 33 | 2 | NA | Fig. 2B |
| **XZ1516-QX1211_self** | XZ1516/QX1211 self | 78 | 60 | 18 | 0 | NA | Fig. 2B |
| **XZ1516-QX1211_self** | XZ1516/QX1211 self | 90 | 67 | 23 | 0 | NA | Fig. 2B |
| **XZ1516-QX1211_self** | XZ1516/QX1211 self | 87 | 69 | 18 | 0 | NA | Fig. 2B |
| **XZ1516-QX1211_self** | XZ1516/QX1211 self | 100 | 73 | 27 | 0 | NA | Fig. 2B |
| **QX2511-QX1211_self** | XZ1516 Δtmrl-1/QX1211 self | 76 | 74 | 0 | 2 | 0 | Fig. 2B |
| **QX2511-QX1211_self** | XZ1516 Δtmrl-1/QX1211 self | 51 | 47 | 0 | 4 | 0 | Fig. 2B |
| **QX2511-QX1211_self** | XZ1516 Δtmrl-1/QX1211 self | 64 | 61 | 0 | 3 | 1 | Fig. 2B |
| **QX2518-DL238_self** | XZ1516 with amrl-1 array/DL238 self | 128 | | 4 | 0 | 0 | Fig. 2B |
| **QX2518-DL238_self** | XZ1516 with amrl-1 array/DL238 self | 107 | | 4 | 0 | 0 | Fig. 2B |
| **QX2518-DL238_self** | XZ1516 with amrl-1 array/DL238 self | 135 | | 3 | 0 | 0 | Fig. 2B |
| **QX2519-DL238_self** | XZ1516 with amrl-1 array/DL238 self | 246 | | 17 | 0 | 0 | Fig. 2B |
| **QX2519-DL238_self** | XZ1516 with amrl-1 array/DL238 self | 122 | | 2 | 0 | 0 | Fig. 2B |
| **QX2516-XZ1516_self** | XZ1516 Δtmrl-1Δamrl-1/DL238 self | 152 | 112 | 37 | 3 | | Fig. 2B |
| **QX2515-NIC195_self** | XZ1516(fluorescent males)/NIC195 self | 78 | 56 | 3 | 0 | 19 | Fig. 3C |
| **QX2515-NIC195_self** | XZ1516(fluorescent males)/NIC195 self | 136 | 96 | 3 | 0 | 37 | Fig. 3C |
| **QX2515-QX2530_self** | XZ1516(fluorescent males)/NIC195Δamrl-1 self | 115 | 60 | 25 | 0 | 30 | Fig. 3C |
| **QX2515-QX2530_self** | XZ1516(fluorescent males)/NIC195Δamrl-1 self | 107 | 64 | 25 | 0 | 18 | Fig. 3C |

### Table S5

| Purpose | Fwd name | Fwd primer | Rev name | Rev primer | Strains genotyped |
|---|---|---|---|---|---|
| **verify fog-2 edit** | oZ7 | CGCCGGATTTCAACCAATCGAAAC | oZ8 | CGCCGGATTTCAACCAATCGAAAC | QX2538 and QX2539 |
| **Genotype QX1211 deletion at V:17789593-17789680 to find recombinations in QX1211/XZ1516 F2s** | oZ50 | TGTCCTTGCCATTGTCCCTC | oZ51 | AAGTCGCAGGTCACACTGAG | QX2500 |
| **Genotype QX1211 deletion at V:18291040-18291112 to find recombinations in QX1211/XZ1516 F2s** | oZ52 | TCCAGTGATGTCCAGACAAAAC | oZ53 | CCTTCAAGAACACCGTCGTG | QX2500 |
| **Genotype XZ1516 deletion at V:20751328 (N2)** | oZ64 | GGGTGCGACTACTTAACCGG | oZ65 | CAATGTCCATTGGTCCGTTGG | QX2501 and QX2506 |
| **Genotype QX1211 deletion at V:20031932 (N2 genome coordinates)** | oZ66 | TGCGCTATTGGGACACATTTC | oZ67 | GGTGATGAGCAACACTGTGC | QX2501 and QX2506 |
| **Verify cas9 induced edit of qq200/201** | oZ152 | CGGCTTCTATCGAATTTGGGCAG | oZ154 & oZ129 | CCACCCAAATTGGGAGGATACAAC (external) CGGCTAACACAACTTTTTTGGGATCC (internal) | QX2509 and QX2510 |
| **Verify cas9 induced edit of qq202/203** | oZ121 | GAGAGAGTGTAAGGCCAAACTGATTC | oZ135 & oZ120 | CGGAATTTCTCTCAGCAGACCATC (external) CCAACGGACGTATCACATTCACC (internal) | QX2511 and QX2512 |
| **Verify cas9 induced edit of qq204/205** | oZ121 | GAGAGAGTGTAAGGCCAAACTGATTC | oZ140 & oZ120 | CCATATTGTCTAGAGCTGGAGCAG (external) CCAACGGACGTATCACATTCACC (internal) | QX2513 and QX2514 |

| | | | | | |
|---|---|---|---|---|---|
| **Verify cas9 induced edit of qq208/209** | oZ157 | CTCAAAAACTTTTACGTCAATTTTAGGG | oZ159 | GAAGCGCGTGGAATAGAATGTCAG (external) | QX2516 and QX2517 |
| **Verify cas9 induced edit of qq208/209** | oZ80 | CACCAATATGAGACATATGAAGAGGGACTTG | oZ82 & oZ86 | CCTGGATGCTTCTCGCTTTTGTGC and CGCATAGATATGGGTGTCAACAAATGATTCAG | QX2516 and QX2517 |
| **Verify cas9 induced deletion of qq210** | oZ146 | GTTGATGATAAATAAGGTCTTGCTCC | oZ145 & oZ147 | CAGAGATTTCTACACTAAATTCCCGGG and GCACAGCACTGGTTTACGAGTCTC | QX2530 |
| **Verify cas9 induced edit of qq211** | oZ249 | CGATGACGATCGTTGAAGATAAGTTGAG | oZ248 & oZ250 | GGAAATCGTACTCCCATTCATCGTATCTC and GCAGTTATGGCTCAACATTCAAACCATC | QX2537 |
| **Amplify amrl-1 CDS with homology to pWM1 (generate pWM4)** | oZ164 | GTTGGGAAACACTTTGCTCAcaagttATGACGTCGTTTATGATAAACAAGAAGCAAC | oZ165 | GAATGCTTGAAAGGATTTTGCATTTATTTAGACAATCTCAAAAGAGGAGTCAGTG | NA |
| **Amplify amrl-1 promoter & CDS with homology to pWM1 (generate pWM3)** | oZ162 | CACGACGTTGTAAAACGACGGCCAGTGGTTACAAAAGAAAGCCAGTTTCATCG | oZ165 | GAATGCTTGAAAGGATTTTGCATTTATTTAGACAATCTCAAAAGAGGAGTCAGTG | NA |
| **Amplify tmrl-1.a with homology to pWM8 (generate pWM11)** | oZ203 | GTCGACGGTAccggtagaaaaacattcgtaAGAGGTGCTTCGGACGTCGC | oZ204 | ttggtaatggtagcgaccggcgctcagttgTTATTCGAGTGGACTGGTTTTGGTAGC | NA |
| **Amplify tmrl-1.b with homology to pWM8 (generate pWM12)** | oZ205 | GTCGACGGTAccggtagaaaaacattcgtaTTTCAGGTGATGCACGGGGTTGG | oZ204 | ttggtaatggtagcgaccggcgctcagttgTTATTCGAGTGGACTGGTTTTGGTAGC | NA |
| **Amplify B0250.8 with homoglogy to pWM8 (generate pWM21)** | oZ217 | GTCGACGGTAccggtagaaaaacattcgtaATGCCAGCACAAAAAGGAGTACAAG | oZ218 | ttggtaatggtagcgaccggcgctcagttgTTATACAGAAGAAGTGGTGTTGGAAACGAG | NA |
| **Amplify amrl-1 CDS with homology to pWM8 (generate pWM25)** | oZ224 | gtcgacggtaccggtagaaaaacattcgtagATGACGTCGTTTATGATAAACAAGAAGC | oZ225 | ggtagcgaccggcgctcagttggaattTTAGACAATCTCAAAAGAGGAGTCAGTG | NA |

| | | | | | |
|---|---|---|---|---|---|
| **Introduce stop codon in tmrl-1 in pWM11 to generate pWM17 (using NEB site-directed mutagenesis)** | oZ288 | gcacaactccgtagaatcctctgctc | oZ289 | gaaacgtctctacaagtgcttgaacgg | NA |

### Table S6

| Description | gRNA name | gRNA sequence | Repair name | Repair template sequence | Strains generated |
|---|---|---|---|---|---|
| **Generate fog-2(qq212)** | oZ10 | GGCAUGUCAGAGAAUGAUGG | oZ9 | aaactcatagtttttataatttcagctccggttaaccatcattctctgacatgccaatggagacggttta | QX2538 and QX2539 |
| **dpy-10 co-injection marker; used in Cas9 injections** | oZ30 | GCUACCAUAGGCACCACGAG | oZ31 | CACTTGAACTTCAATACGGCAAGATGAGAATGACTGGAAACCGTACCGCATGCGGTGCCTATGGTAGCGGAGCTTCACATGGCTTCAGACCAACAGCCTAT | |
| **generate CINR recombinant** | gSZ71 | TCGAGATGAGAAGATCACGT | NA | NA | QX2506 |
| **target FUN_019827; paired with gSZ86 to generate qq200 and qq201; paired with gSZ83 & gSZ87 to generate qq204 and qq205** | gSZ85 | ACAAGCAGTAATTATTAACG | NA | NA | QX2509, QX2510, QX2513, QX2514 |
| **target FUN_019830; paired with gSZ85 to generate qq200 and qq201** | gSZ86 | CAACTGCTTATTCTGTGCTG | NA | NA | QX2509 and QX2510 |
| **target tmrl-1; paired with gSZ108 to generate qq202 and qq203** | gSZ83 | CCAGACCACTTTGGATCCCG | NA | NA | QX2511, QX2512, QX2513, QX2514 |
| **target tmrl-1; paired with gSZ83 to generate qq202 and qq203** | gSZ108 | GGCUUCCUGAGAAGGCAUGU | NA | NA | QX2511 and QX2512 |

| | | | | | |
|---|---|---|---|---|---|
| **target FUN_019830; paired with gSZ83 & gSZ85 to generate qq204 and qq205** | gSZ87 | CTGCGAGACCCATGACACGT | NA | NA | QX2513 and QX2514 |
| **target amrl-1; paired with gSZ112 to generate qq208 and qq209** | gSZ111 | CGAUGUGAAAACUGGACAUG | NA | NA | QX2516 and QX2517 |
| **target amrl-1; paired with gSZ111 to generate qq208 and qq209** | gSZ112 | AGCAACUGAAAGAAAAGAUC | NA | NA | QX2516 and QX2517 |
| **target amrl-1; paired with gSZ128 to generate qq210** | gSZ78 | GCATCCAGGATGGGAACACG | NA | NA | QX2530 |
| **target amrl-1; paired with gSZ78 to generate qq210** | gSZ128 | CATAAGAATAAGTatCACCC | NA | NA | QX2530 |
| **target B0250.8; pared with gSZ122 to generate qq211** | gSZ119 | GAACCGTTTTTAAAACCACT | NA | NA | QX2537 |
| **target B0250.8; pared with gSZ122 to generate qq211** | gSZ122 | CTTCTGTATAATTTTCATCC | NA | NA | QX2537 |

### Table S7

| | G(+) | # healthy | # phenoA | # phenoB | # phenoC | # phenoD | # phenoE | # phenoF | | A - messy, exploded, lollipop type shape, very much dead |
|---|---|---|---|---|---|---|---|---|---|---|
| **XZ tet:tmrl-1 repA** | 47 | 2 | 11 | 16 | 1 | 17 | 0 | 0 | | B - older than L2 worm (could be anywhere from L3-YA), straightened out, dead (or very sick) |
| **XZ tet:tmrl-1 repB** | 24 | 5 | 14 | 4 | 1 | 0 | 0 | 0 | | C - folded completely in half |
| **XZ tet:B0250.8** | 54 | 0 | 2 | 0 | 0 | 2 | 40 | 10 | | D - smaller worms, closer to L1-L2, mostly straightened out (some straighter than others), dead (or very sick) |
| **DL tet:B0250.8** | 42 | 0 | 0 | 0 | 0 | 5 | 11 | 26 | | E - L1-2 sluggish |
| **N2 tet:B0250.8** | 51 | 0 | 0 | 0 | 0 | 1 | 25 | 25 | | F - other sick |

### References:

1. R. W. Beeman, K. S. Friesen, R. E. Denell, Maternal-effect selfish genes in flour beetles. *Science* **256**, 89–92 (1992).

2. E. Ben-David, A. Burga, L. Kruglyak, A maternal-effect selfish genetic element in Caenorhabditis elegans. *Science* **356**, 1051–1055 (2017).

3. H. S. Seidel, M. V. Rockman, L. Kruglyak, Widespread Genetic Incompatibility in \emphC. Elegans Maintained by Balancing Selection. *Science* **319**, 589–594 (2008).

4. E. Ben-David, P. Pliota, S. A. Widen, A. Koreshova, T. Lemus-Vergara, P. Verpukhovskiy, S. Mandali, C. Braendle, A. Burga, L. Kruglyak, Ubiquitous Selfish Toxin-Antidote Elements in Caenorhabditis Species. *Curr. Biol.* **31**, 990–1001.e5 (2021).

5. D. Jurėnas, N. Fraikin, F. Goormaghtigh, L. Van Melderen, Biology and evolution of bacterial toxin–antitoxin systems. *Nat. Rev. Microbiol.* **20**, 335–350 (2022).

6. M. LeRoux, M. T. Laub, Toxin-antitoxin systems as phage defense elements. *Annu. Rev. Microbiol.* **76**, 21–43 (2022).

7. A. J. Shultz, T. B. Sackton, Immune genes are hotspots of shared positive selection across birds and mammals. *Elife* **8** (2019).

8. M. D. Daugherty, H. S. Malik, Rules of engagement: molecular insights from host-virus arms races. *Annu. Rev. Genet.* **46**, 677–700 (2012).

9. H. S. Seidel, M. Ailion, J. Li, A. van Oudenaarden, M. V. Rockman, L. Kruglyak, A novel sperm-delivered toxin causes late-stage embryo lethality and transmission ratio distortion in C. elegans. *PLoS Biol.* **9**, e1001115 (2011).

10. L. M. Noble, J. Yuen, L. Stevens, N. Moya, R. Persaud, M. Moscatelli, J. L. Jackson, G. Zhang, R. Chitrakar, L. R. Baugh, C. Braendle, E. C. Andersen, H. S. Seidel, M. V. Rockman, Selfing is the safest sex for Caenorhabditis tropicalis. *Elife* **10** (2021).

11. M. V. Rockman, Parental-effect gene-drive elements under partial selfing, or why do Caenorhabditis genomes have hyperdivergent regions?, *bioRxiv* (2024)p. 2024.07.23.604817.

12. H. Wang, L. Planche, V. Shchur, R. Nielsen, Selfing Promotes Spread and Introgression of Segregation Distorters in Hermaphroditic Plants. *Mol. Biol. Evol.* **41** (2024).

13. D. Lee, S. Zdraljevic, L. Stevens, Y. Wang, R. E. Tanny, T. A. Crombie, D. E. Cook, A. K. Webster, R. Chirakar, L. R. Baugh, M. G. Sterken, C. Braendle, M.-A. Félix, M. V. Rockman, E. C. Andersen, Balancing selection maintains hyper-divergent haplotypes in Caenorhabditis elegans. *Nat Ecol Evol* **5**, 794–807 (2021).

14. L. Long, W. Xu, F. Valencia, A. B. Paaby, P. T. McGrath, A toxin-antidote selfish element increases fitness of its host. *Elife* **12** (2023).

15. A. Burga, E. Ben-David, T. Lemus Vergara, J. Boocock, L. Kruglyak, Fast genetic mapping of complex traits in C. elegans using millions of individuals in bulk. *Nat. Commun.* **10**, 2680 (2019).

16. C. E. Rocheleau, R. M. Howard, A. P. Goldman, M. L. Volk, L. J. Girard, M. V. Sundaram, A lin-45 raf enhancer screen identifies eor-1, eor-2 and unusual alleles of Ras pathway genes in Caenorhabditis elegans. *Genetics* **161**, 121–131 (2002).

17. M. V. Rockman, L. Kruglyak, Recombinational landscape and population genomics of \emphCaenorhabditis elegans. *PLoS Genet.* **5**, e1000419 (2009).

18. S. Zdraljevic, L. Walter-McNeill, H. Marquez, L. Kruglyak, Heritable Cas9-induced nonhomologous recombination in C. elegans. *microPublication Biology* **2023** (2023).

19. S. Robertson, R. Lin, “Chapter One - The Maternal-to-Zygotic Transition in C. elegans” in *Current Topics in Developmental Biology*, H. D. Lipshitz, Ed. (Academic Press, 2015; https://www.sciencedirect.com/science/article/pii/S0070215315000290)vol. 113, pp. 1–42.

20. D. E. Cook, S. Zdraljevic, J. P. Roberts, E. C. Andersen, CeNDR, the Caenorhabditis elegans natural diversity resource. *Nucleic Acids Res.* **45**, D650–D657 (2017).

21. J. H. Gillespie, C. H. Langley, Are evolutionary rates really variable? *J. Mol. Evol.* **13**, 27–34 (1979).

22. C. G. Thomas, W. Wang, R. Jovelin, R. Ghosh, T. Lomasko, Q. Trinh, L. Kruglyak, L. D. Stein, A. D. Cutter, Full-genome evolutionary histories of selfing, splitting, and selection in Caenorhabditis. *Genome Res.* **25**, 667–678 (2015).

23. A. K. Rogers, C. M. Phillips, A Small-RNA-Mediated Feedback Loop Maintains Proper Levels of 22G-RNAs in C. elegans. *Cell Rep.* **33**, 108279 (2020).

24. C. J. Uebel, D. C. Anderson, L. M. Mandarino, K. I. Manage, S. Aynaszyan, C. M. Phillips, Distinct regions of the intrinsically disordered protein MUT-16 mediate assembly of a small RNA amplification complex and promote phase separation of Mutator foci. *PLoS Genet.* **14**, e1007542 (2018).

25. C. M. Phillips, T. A. Montgomery, P. C. Breen, G. Ruvkun, MUT-16 promotes formation of perinuclear mutator foci required for RNA silencing in the C. elegans germline. *Genes Dev.* **26**, 1433–1444 (2012).

26. K. J. Reed, J. M. Svendsen, K. C. Brown, B. E. Montgomery, T. N. Marks, T. Vijayasarathy, D. M. Parker, E. O. Nishimura, D. L. Updike, T. A. Montgomery, Widespread roles for piRNAs and WAGO-class siRNAs in shaping the germline transcriptome of Caenorhabditis elegans. *Nucleic Acids Res.* **48**, 1811–1827 (2020).

27. A. K. Rogers, C. M. Phillips, Disruption of the mutator complex triggers a low penetrance larval arrest phenotype. *microPublication Biology*, doi: 10.17912/micropub.biology.000252 (2020).

28. E. C. Andersen, T. C. Shimko, J. R. Crissman, R. Ghosh, J. S. Bloom, H. S. Seidel, J. P. Gerke, L. Kruglyak, A Powerful New Quantitative Genetics Platform, Combining \emphCaenorhabditis elegans High-Throughput Fitness Assays with a Large Collection of Recombinant Strains. *G3* **5**, g3.115.017178–920 (2015).

29. Y. V. Makeyeva, M. Shirayama, C. C. Mello, Cues from mRNA splicing prevent default Argonaute silencing in C. elegans. *Dev. Cell* **56**, 2636–2648.e4 (2021).

30. W.-S. Wu, W.-C. Huang, J. S. Brown, D. Zhang, X. Song, H. Chen, S. Tu, Z. Weng, H.-C. Lee,

pirScan: a webserver to predict piRNA targeting sites and to avoid transgene silencing in C. elegans. *Nucleic Acids Res.* **46**, W43–W48 (2018).

31. D. Zhang, S. Tu, M. Stubna, W.-S. Wu, W.-C. Huang, Z. Weng, H.-C. Lee, The piRNA targeting rules and the resistance to piRNA silencing in endogenous genes. *Science* **359**, 587–592 (2018).

32. W. Tang, S. Tu, H.-C. Lee, Z. Weng, C. C. Mello, The RNase PARN-1 Trims piRNA 3' Ends to Promote Transcriptome Surveillance in C. elegans. *Cell* **164**, 974–984 (2016).

33. U. Seroussi, A. Lugowski, L. Wadi, R. X. Lao, A. R. Willis, W. Zhao, A. E. Sundby, A. G. Charlesworth, A. W. Reinke, J. M. Claycomb, A comprehensive survey of C. elegans argonaute proteins reveals organism-wide gene regulatory networks and functions. *Elife* **12** (2023).

34. W. Gu, M. Shirayama, D. Conte Jr, J. Vasale, P. J. Batista, J. M. Claycomb, J. J. Moresco, E. M. Youngman, J. Keys, M. J. Stoltz, C.-C. G. Chen, D. A. Chaves, S. Duan, K. D. Kasschau, N. Fahlgren, J. R. Yates 3rd, S. Mitani, J. C. Carrington, C. C. Mello, Distinct argonaute-mediated 22G-RNA pathways direct genome surveillance in the C. elegans germline. *Mol. Cell* **36**, 231–244 (2009).

35. F. K. Nelson, D. L. Riddle, Functional study of the Caenorhabditis elegans secretory-excretory system using laser microsurgery. *J. Exp. Zool.* **231**, 45–56 (1984).

36. W. C. Forrester, G. Garriga, Genes necessary for C. elegans cell and growth cone migrations. *Development* **124**, 1831–1843 (1997).

37. S. Liégeois, A. Benedetto, G. Michaux, G. Belliard, M. Labouesse, Genes required for osmoregulation and apical secretion in Caenorhabditis elegans. *Genetics* **175**, 709–724 (2007).

38. M. P. Bagijn, L. D. Goldstein, A. Sapetschnig, E.-M. Weick, S. Bouasker, N. J. Lehrbach, M. J. Simard, E. A. Miska, Function, targets, and evolution of Caenorhabditis elegans piRNAs. *Science* **337**, 574–578 (2012).

39. A. Ashe, A. Sapetschnig, E.-M. Weick, J. Mitchell, M. P. Bagijn, A. C. Cording, A.-L. Doebley, L. D. Goldstein, N. J. Lehrbach, J. Le Pen, G. Pintacuda, A. Sakaguchi, P. Sarkies, S. Ahmed, E. A. Miska, piRNAs can trigger a multigenerational epigenetic memory in the germline of C. elegans. *Cell* **150**, 88–99 (2012).

40. H.-C. Lee, W. Gu, M. Shirayama, E. Youngman, D. Conte Jr, C. C. Mello, C. elegans piRNAs mediate the genome-wide surveillance of germline transcripts. *Cell* **150**, 78–87 (2012).

41. M. Shirayama, M. Seth, H.-C. Lee, W. Gu, T. Ishidate, D. Conte Jr, C. C. Mello, piRNAs initiate an epigenetic memory of nonself RNA in the C. elegans germline. *Cell* **150**, 65–77 (2012).

42. A. E. Sundby, R. I. Molnar, J. M. Claycomb, Connecting the Dots: Linking Caenorhabditis elegans Small RNA Pathways and Germ Granules. *Trends Cell Biol.* **31**, 387–401 (2021).

43. J. Vedanayagam, Small RNA-mediated suppression of sex chromosome meiotic conflicts during Drosophila male gametogenesis. *Biochem. Soc. Trans.* **53**, 281–291 (2025).

44. N. Phadnis, H. A. Orr, A single gene causes both male sterility and segregation distortion in Drosophila hybrids. *Science* **323**, 376–379 (2009).

45. J. Bladen, H.-J. Nam, N. Phadnis, Transformation of meiotic drive into hybrid sterility in Drosophila.

*Genetics* **228**, iyae133 (2024).

46. E. C. Andersen, J. S. Bloom, J. P. Gerke, L. Kruglyak, A variant in the neuropeptide receptor npr-1 is a major determinant of \emphCaenorhabditis elegans growth and physiology. **10**, e1004156 (2014).

47. E. Ben-David, J. Boocock, L. Guo, S. Zdraljevic, J. S. Bloom, L. Kruglyak, Whole-organism eQTL mapping at cellular resolution with single-cell sequencing. *Elife* **10** (2021).

48. A. M. Bhagwat, J. Graumann, R. Wiegandt, M. Bentsen, J. Welker, C. Kuenne, J. Preussner, T. Braun, M. Looso, multicrispr: gRNA design for prime editing and parallel targeting of thousands of targets. *Life Sci Alliance* **3** (2020).

49. R. Core, TEAM, 2017. R: A language and environment for statistical computing. R Foundation for Statistical Computing, Vienna, Austria. *Online: https://www. r-project. org* (2022).

50. H. Li, Minimap2: pairwise alignment for nucleotide sequences. *Bioinformatics* **34**, 3094–3100 (2018).

51. S. Mao, Y. Qi, H. Zhu, X. Huang, Y. Zou, T. Chi, A Tet/Q Hybrid System for Robust and Versatile Control of Transgene Expression in C. elegans. *iScience* **11**, 224–237 (2019).

52. H.-G. Drost, A. Gabel, I. Grosse, M. Quint, Evidence for active maintenance of phylotranscriptomic hourglass patterns in animal and plant embryogenesis. *Mol. Biol. Evol.* **32**, 1221–1231 (2015).

53. S. Zdraljevic, C. Strand, H. S. Seidel, D. E. Cook, J. G. Doench, E. C. Andersen, Natural variation in a single amino acid substitution underlies physiological responses to topoisomerase II poisons. *PLoS Genet.* **13**, e1006891 (2017).

54. W. A. Boyd, M. V. Smith, J. H. Freedman, Caenorhabditis elegans as a model in developmental toxicology. *Methods Mol. Biol.* **889**, 15–24 (2012).

55. T. C. Shimko, E. C. Andersen, COPASutils: an R package for reading, processing, and visualizing data from COPAS large-particle flow cytometers. *PLoS One* **9**, e111090 (2014).
